# Atomistic Mechanism of Calcium-Mediated Inward Rectification of the MthK Potassium Channel by Solid-State NMR and MD Simulations

**DOI:** 10.1101/2025.08.14.670285

**Authors:** Carl Öster, Reinier de Vries, Juan Li, Denis Qoraj, Sascha Lange, Chaowei Shi, Wojciech Kopec, Bert L de Groot, Adam Lange

**Author notes:** These authors contributed equally to this work (Carl Öster, Reinier de Vries). Correspondence should be addressed to W.K., B.L.d.G., or A.L.

## Abstract

Inward rectification is a fundamental but poorly understood phenomenon in potassium channel physiology. Despite its physiological importance, the exact mechanism has remained elusive. In this work, we uncover a previously unrecognized calcium-mediated gating mechanism in the MthK potassium channel that sheds new light on this essential process. By combining state-of-the-art proton-detected solid-state NMR spectroscopy with atomistic molecular dynamics simulations, we reveal that divalent calcium ions bind to a novel site just below the selectivity filter, physically obstructing the outward flow of potassium ions whereas inward flow is still possible - analogous to a molecular ball check valve. Secondly, the binding of Ca^2+^ to the newly identified site leads to stabilization of the selectivity filter and allows us to directly observe ion–ion interactions in the filter. These results offer direct experimental support for the long-debated “direct knock-on” mechanism, in which potassium ions move through the filter, without water co-transport.

## Introduction

Potassium (K^+^) channels are essential membrane proteins that enable rapid and selective permeation of K^+^ ions across cellular membranes. A key structural feature of these channels is the selectivity filter (SF), which coordinates K^+^ ions in four binding sites, S1-S4 (see Fig. 1A), and allows high conductance (>100 pS) while maintaining strong selectivity (ca 100-1000 times) compared to similar, but smaller, sodium (Na^+^) ions^1,2^ which are characterized by a higher desolvation energy upon permeation. The precise mechanism by which ions permeate through the SF remains debated^3^, with two prevailing models: the soft knock-on mechanism, in which ions and water alternate^4^, and the direct knock-on mechanism, which involves direct ion-ion contacts and excludes water from the SF^5,6^.

**Figure 1.**
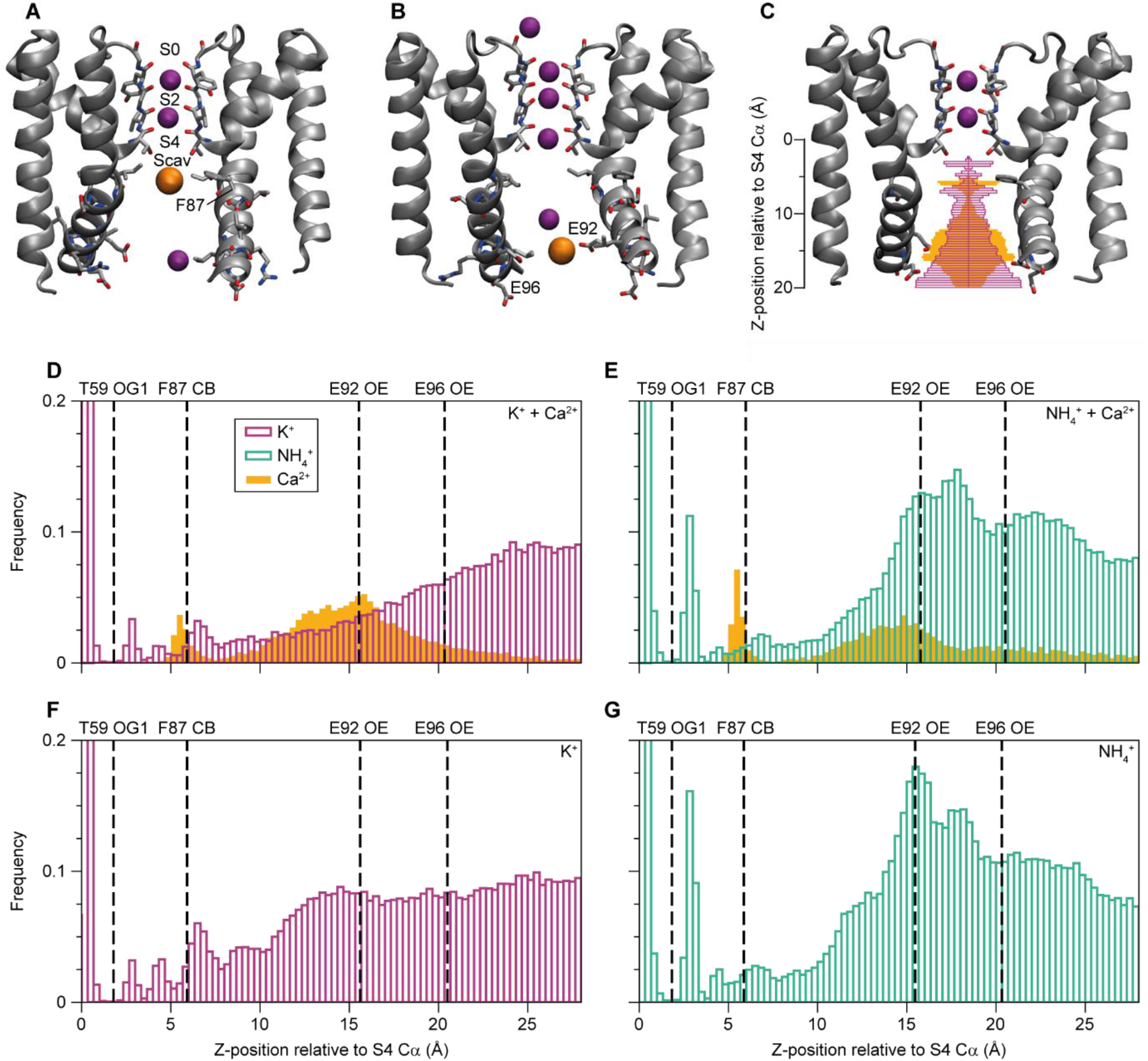
Calcium (Ca^2+^) binding in the MthK cavity under voltage. (A) Representative snapshot of a Ca^2+^ ion bound below the SF under positive voltage. (B) Representative snapshot of a Ca^2+^ ion bound at the glutamate ring under positive voltage (C) Ion densities in the cavity relative to T59 from MD simulations under +300 mV over 10 replicates with a KCl + CaCl_2_ mixture mapped onto the structure of MthK pore domain. (D-G) Ion densities in the cavity relative to T59 from MD simulations under +300 mV over 10 replicates (D) KCl + CaCl_2_, (E) NH_4_Cl + CaCl_2_, (F) KCl, (G) NH_4_Cl.

K^+^ channels can be further divided into different classes. One of these classes is the group of calcium (Ca^2+^)-activated K^+^ channels. Most of these channels open in the presence of calcium ions. Large-conductance K^+^ channels (BK channels) play key roles in muscle contraction, neurotransmission, and blood pressure regulation and display very high conductance rates, even by K^+^ channel standards^7–9^. The archaeal channel MthK serves as a model system for Ca^2+^-activated K^+^ channels. Similarly to BK channels, MthK has a transmembrane pore and a cytoplasmic RCK ring that binds intracellular Ca^2+^ to promote channel opening^10–15^. However, it lacks the voltage sensing domain (VSD) found in eukaryotic Ca^2+^-activated K^+^ channels. Interestingly, Ca^2+^ may also play a secondary role in the ion conductance mechanism by interacting directly with the pore domain, even in the absence of the RCK ring. These interactions result in Ca^2+^-dependent inward rectification, attributed in part to two sets of glutamate residues near the pore entrance (E92 and E96 in MthK)^16,17^. Inward rectification depending on divalent cations is a common effect in K^+^ selective channels. Inwardly rectifying channels, so called Kir channels, are blocked by Mg^2+^ ions and polyvalent cations (e.g. polyamines) in a voltage dependant fashion^18–21^. It has also been shown that BK channels can be blocked by divalent cations such as Mg^2+^ and Ca^2+ 22^. However, the mechanism by which these ions interact with and block the pore is poorly understood. It has been previously proposed that Ca^2+^ and polyamines both block MthK by entering the S4 K^+^ ion binding site^23,24^.

MthK is an attractive model system since there are several structures available of open, closed and inactivated states of the full length channel as well as the pore domain, which due to its small size is especially attractive for MD simulations^13,24,25^. The size and homotetrameric arrangement also makes the MthK pore domain suitable for solid-state NMR experiments. Solid-state NMR has been applied in studies of several different K^+^ channels to gain understanding of gating and ion conduction mechanisms at atomic resolution and under native-like conditions, often in combination with MD simulations^26–32^.

In this study, we use a combination of solid-state NMR and atomistic molecular dynamics (MD) simulations to investigate how Ca^2+^ influences ion occupancy in the SF of MthK. Our findings reveal two distinct Ca^2+^ binding sites in the pore domain. In addition to the previously described glutamate ring, we find that Ca^2+^ can bind in proximity to a phenylalanine below the SF. Ca^2+^ binding blocks K^+^ permeation and alters ion occupancy within the SF, for example by stabilising an otherwise unoccupied SF ion binding site. Both experimental and computational data support a water-free, direct knock-on mechanism of ion permeation, consistent with observations from other K+ channels^31,33^. These results provide new insights into the interplay between Ca^2+^ binding and ion conduction in MthK.

## Results and Discussion

### Ca^2+^ binding sites below the SF identified in MD simulations

We performed separate MD simulations using either KCl or NH_4_Cl, with and without the addition of CaCl_2_. This represents the different sample conditions used in our solid-state NMR experiments. Simulations of the MthK pore under positive voltage, resulting in outward permeation, reveal two main Ca^2+^ binding sites. Figure 1A shows the upper Ca^2+^ binding site close to F87 in transmembrane helix 2 (TMH2), just below the cavity K^+^ binding site (Scav). The second binding site is found near the intracellular pore entrance formed by a ring of glutamate residues (E92, Fig. 1B). A second glutamate ring (E96, further away from the SF) does, however, not show much Ca^2+^ binding, possibly because the distance between the sidechains of E96 is too large for opposing sidechains to coordinate a Ca^2+^ ion. The ion densities observed in the simulations for K^+^ (purple) and Ca^2+^ (orange) are plotted onto the structure of the MthK pore domain in Fig. 1C. The same data is plotted as a bar plot in Fig. 1D. The Ca^2+^ density around F87 is sharp in simulations with both K^+^ (Fig. 1C and D) and ammonium (NH_4_^+^) (Fig. 1E), while the Ca^2+^ density around E92 is broader. The Ca^2+^ occupancy is higher at the E92 binding site compared to the site near F87 in simulations with K^+^. The Ca^2+^ density peak at the glutamate ring is broader, since the four negatively charged glutamate residues allow for different ion binding configurations, whereas the F87 site only allows a single Ca^2+^ position. Compared to simulations without Ca^2+^ (Fig. 1 F and G), the K^+^ and NH_4_^+^ densities at both sites are also reduced in the presence of Ca^2+^, especially near the glutamate ring (Fig. 1C to G). Comparing simulations with K^+^ and NH_4_^+^ reveals a qualitatively similar picture. Increased Ca^2+^ density is observed in the same places, but with the balance slightly shifted between the two, likely because of the stronger binding of NH_4_^+^ to the glutamate ring, indicated by a peak near the E92 side-chain oxygen atoms and higher overall densities (Fig.1 E and G). In simulations at negative (i.e. ion flow in the inward direction) or without voltage (Figs. S1 and S2), binding of Ca^2+^ directly below the SF (around F87) is drastically reduced, whereas the Ca^2+^ density at the glutamate ring remains similar compared to the simulations with positive voltage. This could be a partial explanation for increased inward rectification of MthK with higher Ca^2+^ concentration; a mechanism where the permeation pathway is only blocked under positive voltage.

From the simulations with an applied electric field, we also computed the single channel currents with and without the addition of Ca^2+^ ions (Fig. S3). As previously shown, the CHARMM36m force field does not reproduce the single channel conductance and inward rectification of MthK^34^, however, the main effect we are interested in here is the effect of Ca^2+^ on the current. While there is no significant difference in simulations with and without Ca^2+^, there are relatively few permeation events and a large variance. It has previously been shown that helix dynamics affect the permeability of the channel^13^ and the use of restraints can increase permeation^35^. We therefore also performed simulations with position restraints on the backbone atoms of the lower helices (residues 86-98). This results in ~5-fold increase in current under positive voltage. Here, there is a significant reduction due to the addition of Ca^2+^ under positive voltage, while permeation under negative voltage is unaffected, agreeing with previous experimental data^17,36^. The reduction in current is explained by Ca^2+^ blocking the permeation pathway at the F87 site (Supplementary movie 1) with Ca^2+^ binding at the same binding sites compared to the unrestrained simulations. We also performed electrophysiology experiments on our MthK construct, produced in the same way as the NMR samples, which show the same inwardly rectifying behaviour (Fig. S3).

### Effects of Ca^2+^ binding revealed by solid-state NMR

Next, we performed ^1^H-detected solid-state NMR experiments on the MthK pore domain (in the following simply referred to as MthK). This resulted in spectra with a similar pattern as observed before for other ion channels with similar structures (e.g NaK, NaK2K)^31,33^. 2D (H)NH spectra of H_2_O back-exchanged ^2^H^13^C^15^N labelled MthK (Fig. 2A and B) show peaks for solvent exposed residues, including some highly intense peaks and a broad background signal. Chemical shift assignments were performed based on a combination of ^1^H-detected (of H_2_O back-exchanged ^2^H^13^C^15^N labelled MthK) and additionally recorded ^13^C-detected (of ^13^C^15^N labelled MthK) solid-state NMR experiments. Unambiguous assignments were achieved for the upper part of the transmembrane region of the protein, the SF and the loop connecting the SF to TMH2. Additionally, a few residues in TMH2 below the SF could be assigned based on ^1^H detected triple sensitivity-enhanced 4D experiments^37^, despite severe line broadening of the peaks in that region (Fig. S4). Assigned residues (L28-L79 and T86-V89, excluding A58, tables S1 and S2) are labelled with orange and blue in Figure 2C. Orange labels represent residues for which amide protons are assigned based on ^1^H detected spectra of a H_2_O back-exchanged perdeuterated sample (also labelled in Fig. 2A). Blue labels represent residues that could be assigned based on ^13^C and/or ^1^H detected experiments, but where the amide protons were not visible in ^1^H detected experiments. These parts are likely protected from H/D exchange due to their location in the lipid bilayer. In order to detect bound ions in the SF, we use ^15^NH_4_^+^ ions as mimics for K^+^ ions^33,38^. Fig. 2A and B show that 2D (H)NH spectra of samples with ^15^NH_4_Cl and KCl, both with 10 mM CaCl_2_ in the buffer, look almost identical (see also Fig. S5 for an overlay of the spectra). The only noticeable difference between the samples is that 2 different conformations can be detected for the SF residues when ^15^NH_4_Cl is used and 3 different conformations for some of the SF residues when KCl is used (see also tables S2 and S3). Note that in the absence of CaCl_2_, only one conformation of the SF residues can be observed in the sample with ^15^NH_4_Cl and two conformations in the sample with KCl (Fig. S6). Fig. 2D shows strip plots from (H)CANH and (H)CONH spectra of the residues surrounding the S2 ion binding site, in the presence (orange) and absence (green) of Ca^2+^ (both samples contain ^15^NH_4_^+^ ions). Two different conformations of all atoms around the S2 ion binding site (Y62: H, N, CA; G61: H, N, CA, CO; V60: CO) can be observed in the sample with Ca^2+^, but only one conformation in the sample without Ca^2+^. As will become clear in the next section, where we describe how the ^15^NH_4_^+^ ion binding pattern in the SF depends on the presence of Ca^2+^ ions in the sample, the different conformations (“A” and “B”) correspond to whether an ion is bound in the S2 ion binding site or not.

**Figure 2.**
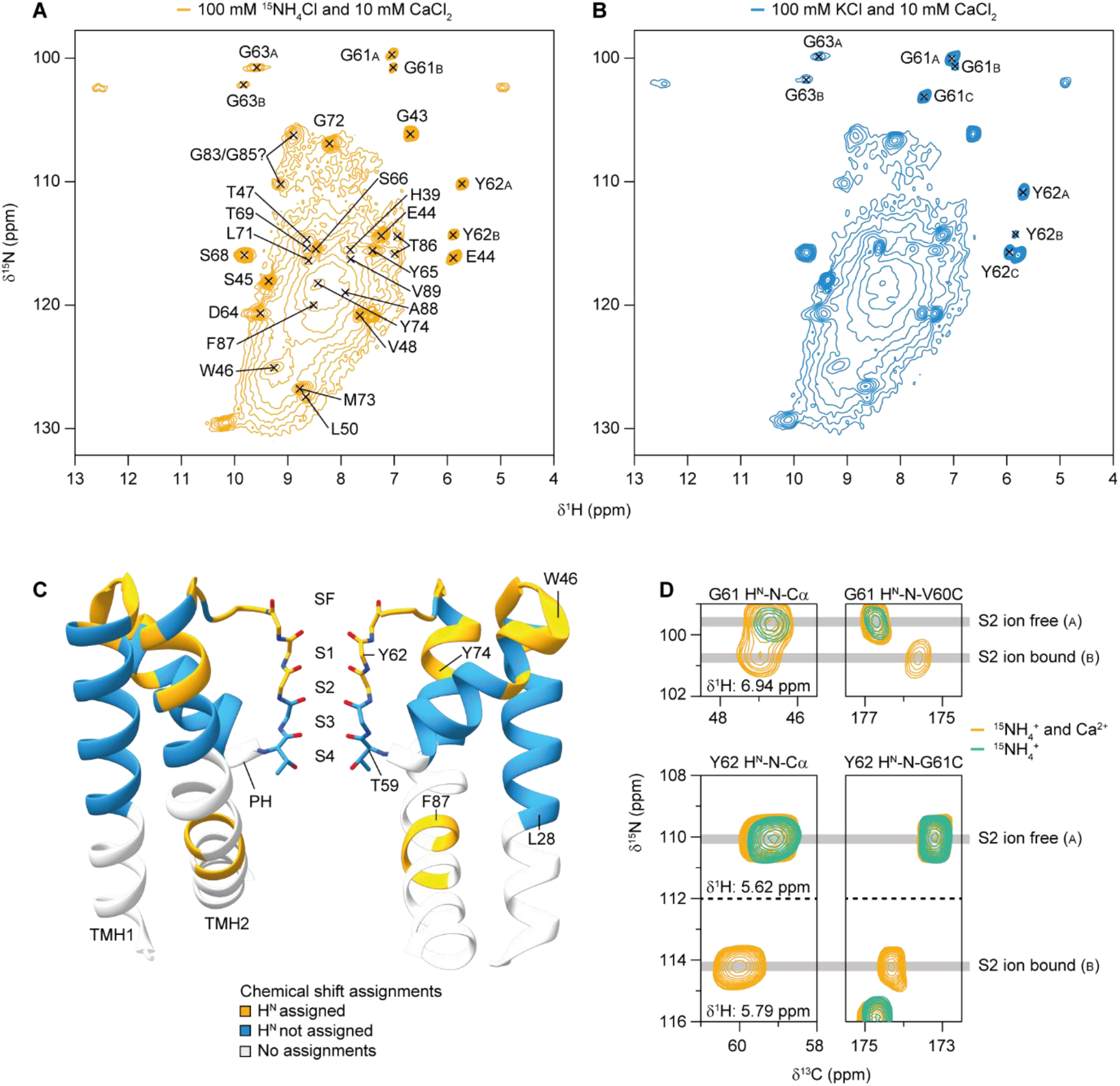
Solid-state NMR analysis of MthK pore domain. 2D (H)NH spectra of (A) MthK with ^15^NH_4_^+^ and Ca^2+^ ions and (B) with K^+^ and Ca^2+^ ions. Residues for which the H^N^ and N chemical shifts are assigned are indicated in (A), only the assigned SF residues are labelled in (B). (C) Chemical shift assignments plotted as a function of residue on the crystal structure of the MthK pore domain (PDB ID: 3LDC). Residues are labelled in orange if the corresponding H^N^ is assigned, in blue if the H^N^ is not assigned and in light grey if no assignments are available. SF = selectivity filter, PH = pore helix, TMH = transmembrane helix. (D) Effects of Ca^2+^ ions on the SF of MthK. The spectra on the left ((H)CANH) show peaks corresponding to H, N and Cα of G61 (top) and Y62 (bottom) with (orange) and without (green) 10 mM CaCl_2_ in the sample buffer. The spectra on the right show peaks corresponding to H^N^ and N of G61 connected to C (CO) of V60 (top) and H^N^ and N of Y62 connected to C (CO) of G61 (bottom).

Another effect of Ca^2+^ ions is increased rigidity, as evident from increased sensitivity in cross-polarization based experiments and the appearance of strong CB-N-H peaks for serine residues in (H)CANH spectra (Fig. S6). The most likely explanation for this effect is that Ca^2+^ binding below the SF reduces the structural heterogeneity caused by conformational dynamics of the lower parts of the transmembrane helices. The heterogeneity of this region leads to a broad background signal for solvent exposed residues, as observed in 2D (H)NH spectra (Fig. 2A and B). Due to this behaviour, it is challenging to obtain unambiguous assignments of residues in the helices below the SF. However, when Ca^2+^ ions are present in the sample it is possible to assign a stretch of residues in TMH2 below the SF (T86-V89, see Fig. S4). Interestingly, this is one of the regions where Ca^2+^ ions are binding according to the MD simulations (Fig. 1A). Additionally, a few intense peaks corresponding to glycines, alanines and leucines could be identified but not unambiguously assigned. These peaks likely correspond to other residues in TMH2 that are slightly stabilized by Ca^2+^ ions but still exist in multiple conformations, as evident by strong line broadening. RMSD (Fig. S7) and RMSF (Fig. S8) analysis of the MD simulations show largest structural deviations compared to the starting structure and strongest dynamics during the simulations for the lower parts of TMH1 and TMH2. There are, however, no differences between simulations with and without Ca^2+^, supporting the hypothesis that the increased sensitivity observed in the solid-state NMR spectra with Ca^2+^ is caused by a reduction of slow conformational dynamics at a timescale of μs to ms that is slower than what can be observed by simulations (a few μs). It has previously been shown that conductance of K^+^ channels depends on the degree of opening of the lower gate (around F97, at the bottom of TMH2), and that opening of the lower gate is coupled to the SF^13^. We simulated how Ca^2+^ affects the opening of the lower gate by performing MD simulations starting from different degrees of opening, with and without the addition of Ca^2+^ ions. The average distance between opposing F97 Cα atoms are shorter in the presence of Ca^2+^ ions (Figs S9 and S10). This effect potentially contributes to the increased rigidity of the pore and the lower K^+^ conduction in the presence of Ca^2+^ ions.

### Additional insight from simulations of the F87A mutant

Since our MD simulations and solid-state NMR experiments showed that Ca^2+^ binding around F87 is responsible for blocking outward permeation in MthK, we performed additional simulations of a F87A mutant to find out whether this residue is essential for Ca^2+^ binding. In simulations under voltage we find that removing this large bulky hydrophobic group in the cavity results in an increase in current. However, the F87A mutant also show inward rectification in the presence of Ca^2+^ ions in both free and restrained simulations (Fig S11). Inspection of the ion densities in the cavity (Fig. S12) show that Ca^2+^ is actually slightly increased in simulations of the F87A mutant compared to in the WT, but the overall ion distributions are very similar. We therefore conclude that F87 is not required for Ca^2+^ binding below Scav, but rather that Ca^2+^ limits the available sidechain conformations of F87.

### ^15^NH_4_^+^ ion binding and ion-ion interactions

We have previously shown that ^1^H detected solid-state NMR experiments can be used to characterize ion binding in K^+^ selective ion channels^33^. This approach takes advantage of the readily detectable ^1^H and ^15^N nuclei of ^15^NH_4_^+^ ions that are suitable mimics for K^+^ ions (see also Fig. 2). Here we apply this method to elucidate the effects interactions between Ca^2+^ ions and the pore domain of MthK have on ^15^NH_4_^+^ binding in the SF. Figure 3 shows assigned ^1^H detected spectra of bound ^15^NH_4_^+^ ions in the absence (Fig. 3A, green spectrum) and presence (Fig. 3B, orange spectrum) of Ca^2+^ ions (assignments in table S4). In the sample without Ca^2+^ ions (Fig. 3A), bound ^15^NH_4_^+^ ions are detected in the S1, S3 and S4 ion binding sites. In the presence of Ca^2+^ ions (Fig. 3B), ^15^NH_4_^+^ ions are detected in all binding sites (S1 to S4). Interestingly, all the peaks resulting from bound ^15^NH_4_^+^ ions in the sample without Ca^2+^ (except for the unassigned peak) can be detected in the sample with Ca^2+^ present. The additional peaks appearing when Ca^2+^ is present in the sample correspond to an ion bound in the S2 ion binding site and second conformations for ions bound in the S1 and S3 ion binding sites (S1_B_ and S3_B_). All the expected cross-peaks between the ^1^H atoms of the ^15^NH_4_^+^ ions and the backbone carbonyl carbons can be detected in cross-polarization based 2D (H)COH spectra. For the sample without Ca^2+^ (Fig. 3A) these peaks are S1_A_H-G61_A_CO, S1_A_H-Y62_A_CO, S3_A_H-T59CO, S3_A_H-V60_A_CO, and S4H-T59CO. All of those peaks are also present in the sample with Ca^2+^(Fig. 3B), with additional peaks confirming the second conformations of S1 and S3 and an ion occupying the S2 ion binding site. The ion bound in S2 only shows cross-peaks to conformation “B” of the SF residues. In the sample with Ca^2+^ ions, cross-peaks for S1_A_H-G63_A_CO and S1_A_H-V60_A_CO can also be detected, most likely due to increased sensitivity in the Ca^2+^ containing sample. (see also Fig. S13 for connections between ^15^NH_4_^+^ ions and aliphatic carbons in the SF). The unassigned ^15^NH_4_ peak in Fig. 3A could either correspond to an ion bound to the glutamate ring below the SF, that showed high occupancy in MD simulations (Fig. 1), or a different conformation of the S3 ion binding site. It is unlikely that it corresponds to any of the other ion binding sites, since it should then have resulted in cross-peaks to other backbone carbons in the SF. The ion binding pattern observed in MthK (summarized in Fig. 3C) points towards the possibility that ions can be bound in adjacent ion binding sites in the SF. Figure 3E shows the build-up of magnetization between ions bound in different ion binding sites as well as between ions and bound water molecules for MthK with Ca^2+^ present. It is not possible to separate the ^1^H chemical shifts of S3_A_, S3_B_, and S_2_ in these spectra and additionally the S1_B_ and S4 ^1^H chemical shifts are too similar to be distinguished. It is therefore not possible to obtain site-specific information on the magnetization transfer between the ions. Nonetheless, the experiments clearly show that magnetization builds up faster between ions than between ions and water molecules meaning that the SF is water-free under the given experimental conditions.

**Figure 3.**
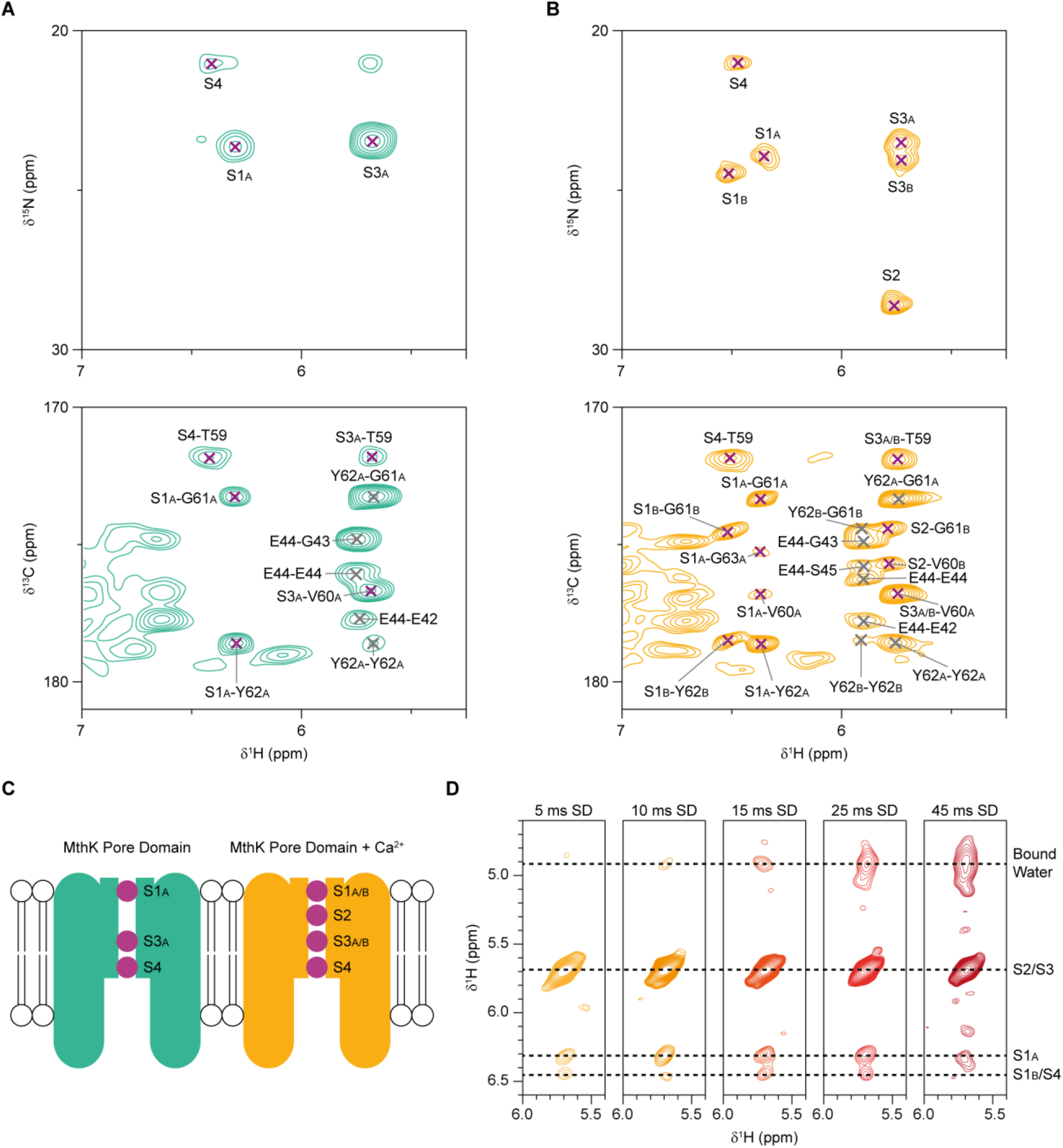
Detection of bound ^15^NH_4_^+^ ions in the MthK pore domain. ^1^H detected INEPT-based 2D hNH spectra of ^15^NH_4_^+^ (top) and CP-based (H)COH spectra (bottom) recorded on ^2^H^13^C^15^N labelled MthK pore domain samples with 100 mM ^15^NH_4_Cl (A, green spectra) and 100 mM ^15^NH_4_Cl + 10 mM CaCl_2_ (B, orange spectra). Peaks involving ^15^NH_4_^+^ ions are labelled with purple crosses and peaks between backbone atoms are labelled with dark grey crosses. The labels in the (H)COH spectra corresponds to ^1^H atoms (left) and carbonyl carbons (right). (C) Scheme of the detected ions in MthK pore domain without (green) and with (orange) CaCl_2_. (D) 2D H(HN)H spectra with varying ^1^H-^1^H spin diffusion mixing times of ^15^NH_4_^+^ ions bound in the SF of MthK, recorded on the sample with 10 mM CaCl_2_.

To get a better view on which ions are simultaneously occupying the SF, we performed additional experiments at a higher external magnetic field strength (1200 MHz ^1^H Larmor frequency, i.e. at 28.2 T) on a sample with a high concentration of Ca^2+^ ions (100 mM). A 3D H(H)NH experiment with 1.16 ms ^1^H-^1^H RFDR mixing reveals several ion-ion contacts (Figure 4). The same ^15^NH_4_^+^ peaks are observed in this spectrum (Fig. 4A) as in the spectrum recorded on a sample with 10 mM Ca^2+^ (Fig. 3B), but a higher Ca^2+^ concentration results in higher sensitivity and the higher magnetic field strength used (1200 MHz vs 600 MHz) further improves sensitivity as well as resolution.

**Figure 4.**
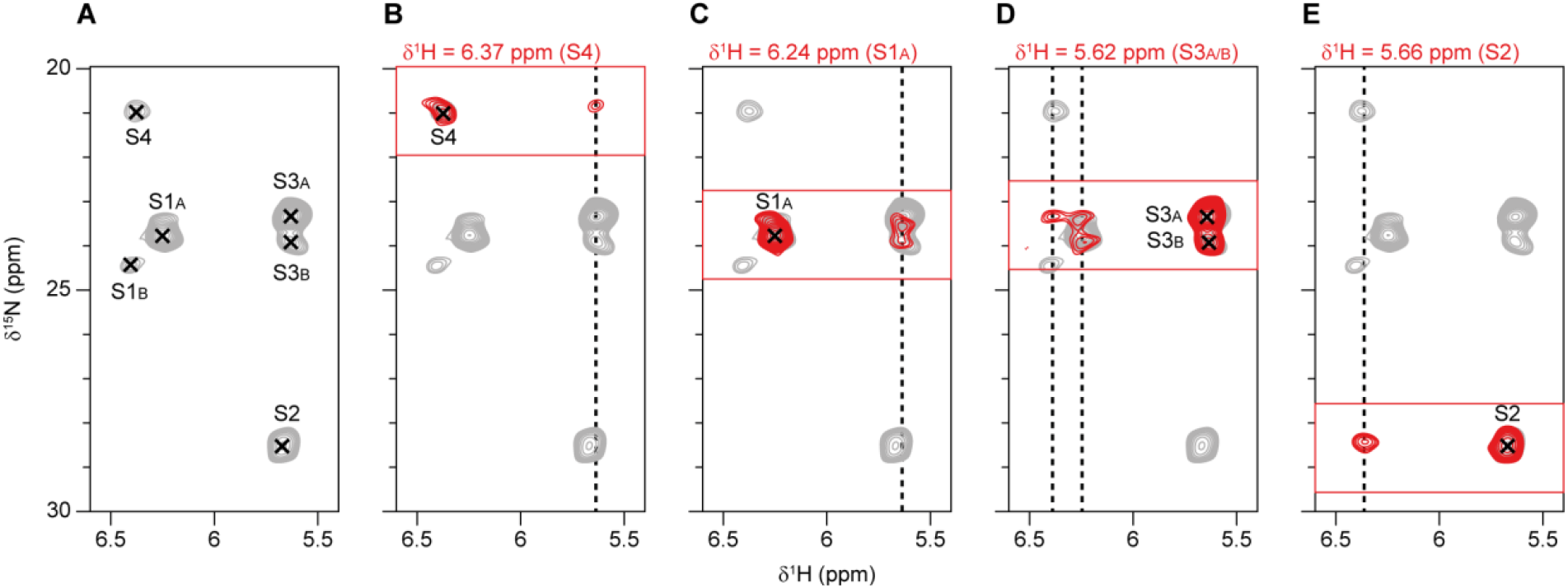
Ion-ion contacts in the MthK pore domain. ^1^H-^15^N 2D projection (grey) of the indirect ^15^N dimension and the direct ^1^H dimension with assignments indicated (A) from a 3D H(H)NH spectrum with 1.16 ms ^1^H-^1^H RFDR mixing. Strip plots (red) taken at the ^1^H chemical shift of the direct dimension for each of the ion peaks showing off-diagonal cross-peaks (B to E: S4, S1A, S3A/B, and S2). The strip plots are overlaid with the ^1^H-^15^N projection in B to E for better visualization of the cross-peaks. The indirect ^1^H chemical shifts of the ion-ion cross-peaks are indicated with dotted lines. The spectrum was recorded on the MthK pore domain with 100 mM ^15^NH_4_Cl and 100 mM CaCl_2_ at 28.2 T and 60 kHz MAS.

Ion-ion contacts between S1_A_ and S3_A/B_ are identified based on cross-peaks from both directions (Fig. 4C and D). The strip plot taken at the ^1^H chemical shift of S3_A_ and S3_B_ (Fig. 4D) clearly shows magnetization transfer from S1_A_ to both S3_A_ and S3_B_. Additionally, transfer to S3_A_ from either S4 or S1_B_ can be seen in Fig. 4D. Magnetization transfer to S4 (Fig. 4B) is coming from S3_A_, S3_B_, or S2, where the ^1^H chemical shift of the cross-peak matches best with S3_A/B_. Fig. 4E shows magnetization transfer to S2, from a peak with a ^1^H chemical shift matching that of an ion bound in the S1_B_ or S4 ion binding site. It is not possible, based on these data, to determine whether there are any cross-peaks between S3_A/B_ and S2 since their ^1^H chemical shifts are too similar. Nonetheless, these data show that it is possible to detect magnetization transfer between ions bound in different ion binding sites and, perhaps most interesting, that the ion bound in the S3 binding site is close in space to (at least) two other ions (most likely ions bound in the S1 and S4 ion binding sites). If we assume that we have a mix of two different ion occupancies, one corresponding to the conformation seen in the sample without Ca^2+^ (S1_A_, S3_A_, and S4, Fig. 3A) and one corresponding to the new peaks appearing when Ca^2+^ is bound below the SF (S1_B_, S2, and S3_B_, Fig. 3B and Fig. 4), then the cross-peaks involving S3_A_ are transfers to S1_A_ and S4, and the cross-peak involving S2 is to S1_B_. This would also agree with the cross-peaks observed between bound ions and backbone carbon atoms in the SF, where S2 shows transfers to G61_B_ and V60_B_ (Fig. 3). However, there is a connection between S1_A_ and S3_B_, which does not agree with a clear distinction between Ca^2+^ free and Ca^2+^ bound SF occupancies. The cross-peaks between S3_A/B_ and the backbone carbon atoms of the SF are only present for conformation “A” of the SF (Fig. 3B, Fig. S13), suggesting that S3_A_ and S3_B_ correspond to slightly different positions within the S3 ion binding site even when no ion is bound in S2. The data at 1200 MHz show that S1_A_ is actually also split in the ^15^N dimension (Fig. 4C), suggesting that an ion bound in the S1 ion binding site also can occupy slightly different positions when no ion is bound in the S2 ion binding site.

### Voltage and Ca^2+^ dependent SF occupancies

Simulations show that the occupancy of K^+^ (and NH_4_^+^) in the SF depends on voltage and whether a Ca^2+^ ion is bound below the SF or not. Figure 5 shows the occupancies based on simulations in the presence of Ca^2+^ ions. The plots are split to show how the K^+^ (Fig. 5A to C) and NH_4_^+^ (Fig. 5D to F) occupancies depend on whether a Ca^2+^ ion is bound below the SF (K^+^ blue bars, NH_4_^+^ orange bars) or not (K^+^ purple bars, NH_4_^+^ green bars). Water molecules (red bars) rarely enter the SF during any of the simulations, in agreement with current (Fig.3) and previous^31,33^ NMR data, as well as previous simulations on other K^+^ selective channels^5,39^, When no Ca^2+^ ion is bound below the SF, the occupancies are similar to simulations without any Ca^2+^ in the simulation box (Fig. S12 and S13). The K^+^ occupancy shifts from mainly KOKO (S1-S4, K = K^+^, O = empty) at negative voltage (Fig. 5A) to KKOK at positive voltage (Fig. 5C), with some K^+^ density observed in S2 at negative and zero voltage and increased density in S4 in simulations at zero voltage compared to negative voltage (but still mainly KOKO, Fig. 5B). With Ca^2+^ bound below the SF the occupancies shift. Under positive voltage the occupancies of S1 and S3 increase, while the occupancies of S2 and S4 decrease when a Ca^2+^ ion is bound below the SF. At zero voltage KKOK becomes the most common state, compared to KOKO without Ca^2+^. Under negative voltage there is very little binding of Ca^2+^ ions below the SF, ions only rarely transiently enter the binding location, so no effect from Ca^2+^ on K^+^ occupancy is observed. Another interesting observation is that the position of an ion in a specific binding site is affected by whether an ion is present in an adjacent ion binding site or not. This is particularly clear when comparing simulations at 0 and 300 mV with and without a Ca^2+^ ion bound below the SF. The position of the K^+^ ion bound in the S1 binding site is shifted downwards when no K^+^ ion is present in the S2 ion binding site (Fig 5B and C). This may potentially explain the nitrogen peak splitting observed in solid-state NMR e.g. for S3_A_ and S3_B_ (Fig. 4B).

**Figure 5.**
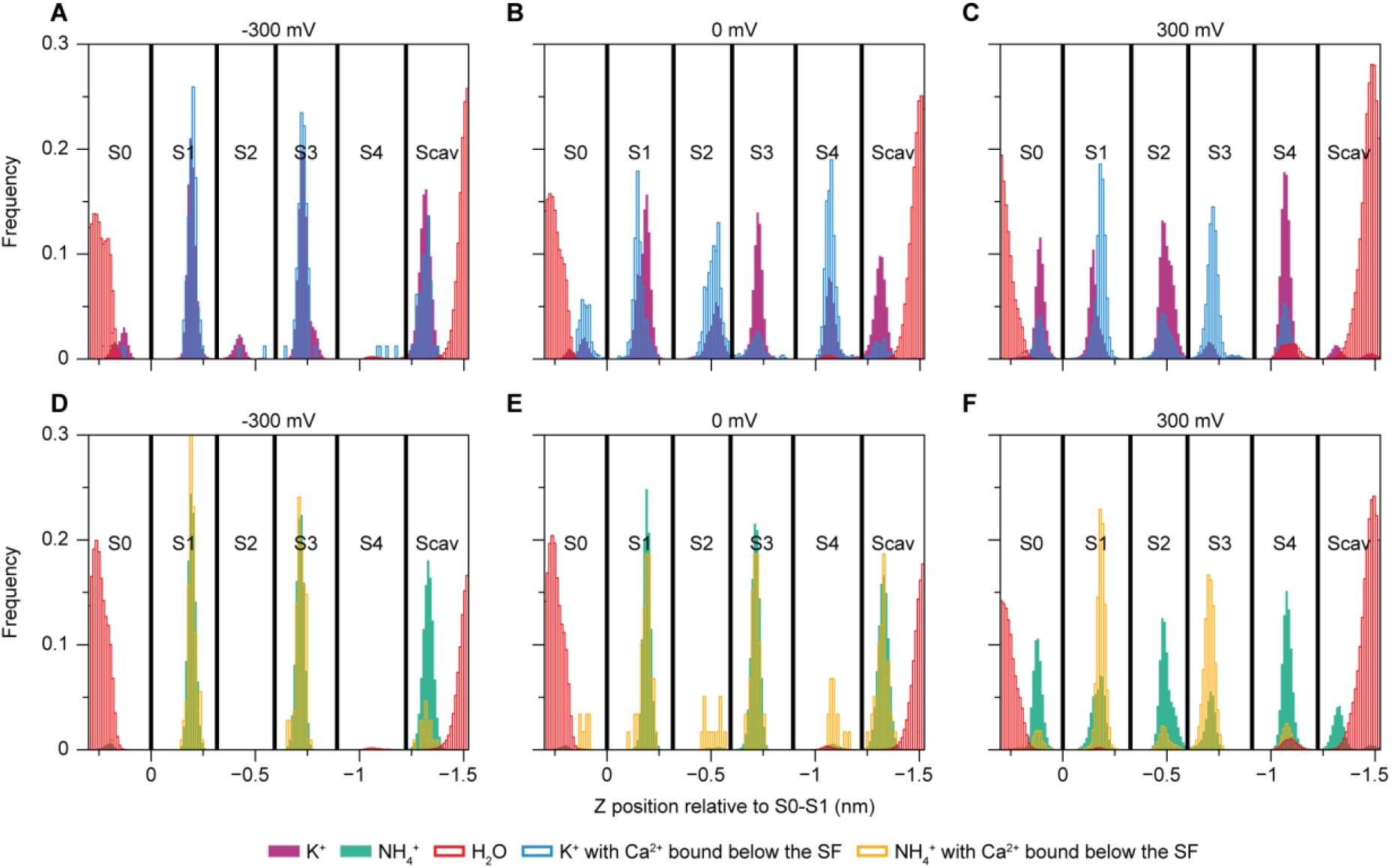
Influence of voltage and Ca^2+^ ions on SF occupancy. (A to C) K^+^ and water occupancies at −300 (A), 0 (B) and +300 (C) mV. (D to F) NH_4_^+^ and water occupancy at −300 (D), 0 (E) and +300 (F) mV. K^+^ and NH_4_^+^ occupancies are sorted and analyzed separately depending on whether a Ca^2+^ ion is bound below the SF (K^+^ in blue, NH_4_^+^ in orange) or not (K^+^ in purple, NH_4_^+^ in green). Water occupancies are shown in red.

Similar effects on occupancies are also observed in simulations where NH_4_^+^ ions are used instead of K^+^ (Fig. 5D to F). Under positive voltage the same shift from KKOK to KOKO is observed when Ca^2+^ is bound below the SF (Fig. 5F). Without positive voltage, Ca^2+^ binding below the SF is even lower than in the case with K^+^ and therefore it is unclear how this would affect the SF NH_4_^+^ occupancy. For both NH_4_^+^ and K^+^ the SF occupancy is voltage dependent and follows a similar trend (Figs 5, S14 and S15). KOKO is dominant under negative voltage, but with increasing positive voltage the occupancies in S2 and S4 increase. While the effect itself is similar, there are some differences between the two cation types. K^+^ exhibits higher S2 and S4 occupancies, even at negative voltages, and the dominant state changes at lower voltage, for example: the occupancy at 300 mV for NH_4_^+^ is very similar to the K^+^ occupancy at 100 mV (Figs. S14 and S15).

## Conclusions

MD simulations reveal two potential Ca^2+^ binding sites below the SF of the MthK pore domain. Effects of Ca^2+^ binding to the pore domain can be observed in solid-state NMR experiments of MthK embedded into a lipid bilayer, as evident by increased rigidity of the protein, changes to the SF conformation and changes to the^15^NH_4_^+^ ion binding pattern upon Ca^2+^ binding below the SF. Ca^2+^ binding allows for chemical shift assignments of a stretch of residues around F87, one of the potential Ca^2+^ binding sites, suggesting reduced conformational dynamics for that region in the presence of Ca^2+^. Ca^2+^ binding to the binding site around F87 is only observed under positive voltage (corresponding to outward flow) in the MD simulations, suggesting that a physical Ca^2+^ block is responsible for the inward rectification of MthK – reminiscent of a ball check valve where Ca^2+^ represents the ball.

The main conclusions of this research are summarized in figure 6. Without Ca^2+^ ions, both inward and outward conduction of K^+^ is unrestricted and the lower part of TMH2 appears to be heterogeneous due to conformational dynamics (Fig. 6A, top). The dominant occupancy of the SF, according to the NMR data of bound ^15^NH_4_^+^ ions, corresponds to ions bound in the S1_A_, S3_A_ and S4 ion binding sites (Fig. 6A, bottom). When Ca^2+^ ions are present, outward K^+^ conduction is blocked by a Ca^2+^ ion bound below the SF close to F87 (Fig. 6B, top). This leads to a stabilization of the pore, especially evident for the lower part of TMH2. Inward flow is not restricted by Ca^2+^ ions and a Ca^2+^ ion bound to the glutamate ring (E92) does not block K^+^ ions from entering or exiting the pore. Based on the different conformations of the SF residues and bound ^15^NH_4_^+^ ions, and the ion-ion contacts detected in NMR experiments at least three different ion occupancy patterns are observed in the SF. We attribute these different SF occupancy patterns to ions bound in S1_A_-S2-S4, S1_A_-S3_B_, and S1_A_-S3_A_-S4 (Fig. 6B bottom).

**Figure 6.**
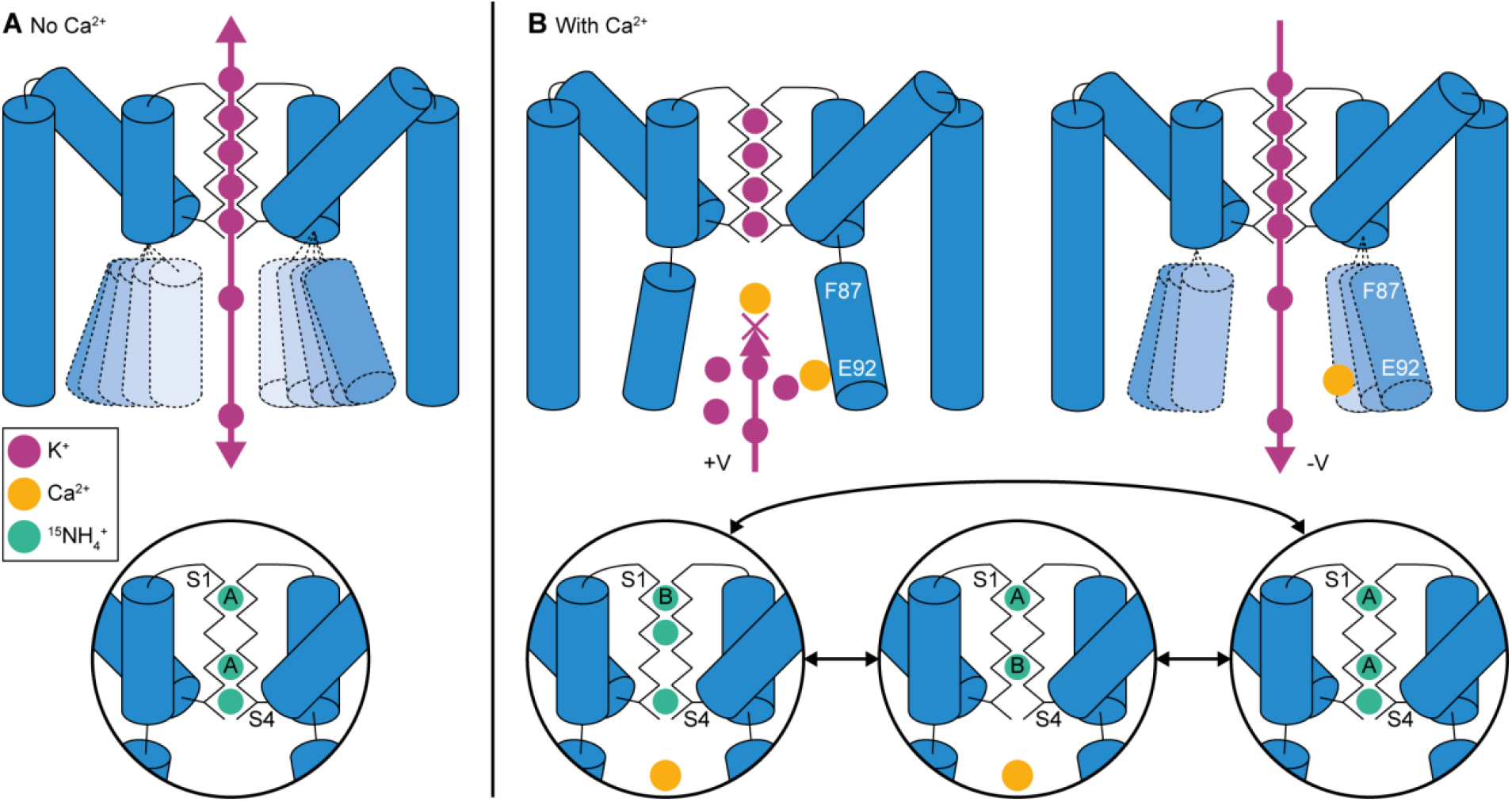
Summary of ball check valve mechanism. (A) Without Ca^2+^ ions, K^+^ ions can flow unhindered inwards and outwards. The lower part of TMH2 undergoes conformational dynamics. The dominant conformation of the SF, as observed in solid-state NMR experiments using ^15^NH_4_^+^ ions, is S1_A_-S3_A_-S4. (B) In the presence of Ca^2+^, the channel is inwardly rectifying. Ca^2+^ ions block outward flow of K^+^ ions leading to a stabilization of TMH2. The dominant configurations of the SF, as observed by solid-state NMR experiments, are S1_B_-S2-S4, S1_A_-S3_B_, S1_A_-S3_A_-S4.

Our MD simulations show that NH_4_^+^ is a good K^+^ mimic to study ion occupancy in MthK. Both K^+^ and NH_4_^+^ show competition in binding to the cavity with Ca^2+^ ions, resulting in the same ion binding sites. They show a similar response to the applied external electric field, shifting the positions of the ions in the SF, although these shifts happen at slightly different voltages. Additionally, they also show the same general shift in occupancy when Ca^2+^ ions bind below the SF, meaning that the presence of a Ca^2+^ ion around F87 has a direct effect on the occupancies of ions in the SF. Solid-state NMR experiments show, similarly to the MD simulations, that the addition of Ca^2+^ leads to different conformations in both samples with K^+^ and ^15^NH_4_^+^. The dominant conformations of the SF appear, however, to be slightly different for K^+^ and ^15^NH_4_^+^.

Last not least, we have detected, for the first time, ion-ion contacts in the SF of a K^+^ channel using solid-state NMR ^1^H-^1^H correlation experiments. The exact occupancies of the different ion binding sites are challenging to determine, but the data show that in the presence of Ca^2+^ multiple configurations can be observed and ^15^NH_4_^+^ ions can occupy adjacent ion binding sites in the SF. Both NMR and MD data show that the exact positions of the ions in the ion binding sites of the SF can vary depending on the occupancies of adjacent ion binding sites, and that water molecules do not regularly enter the central ion binding sites. The simulated SF occupancies for NH_4_^+^ ions show a similar effect as the solid-state NMR data for the S1 and S3 ion binding sites. An opposite effect is observed in simulations compared to NMR experiments for the ion bound in the S2 ion binding site, in simulations the occupancy is higher without Ca^2+^ whereas in NMR experiments S2 is only detected in the presence of Ca^2+^. The combined MD simulation and solid-state NMR approach used here allowed us to shed light on the inwardly rectifying properties of MthK in the presence of Ca^2+^, an interesting feature for a channel that is (in its full-length form with the RCK domains) also activated by Ca^2+^.

## Methods

### MD

The high-resolution crystal structure (PDB: 3LDC)^40^ was mutated back to the WT sequence using the PyMOL mutagenesis tool (H68S C77V). This structure was embedded in a POPC bilayer and a solvated box with ~0.8 mM KCl using the CHARMM-GUI bilayer builder^41^. The box was equilibrated following the 6 step equilibration procedure provided by CHARMM-GUI where position restraints are gradually reduced in consecutive equilibration runs. Starting configurations for the different replicates were generated by running a short (100 ps) NVT simulation. In simulations with NH_4_^+^ ions the same box was used, replacing the K^+^ ions by NH_4_^+^ ions. For simulations with Ca^2+^ we used a multisite Ca^2+^ model^42^, gmx genion was used to add 10 Ca^2+^ ions (~30 mM) and these were converted to the multisite model using the provided script. Systems were simulated using GROMACS 2022.6^43,44^. For voltage simulations an external electric field was applied along the Z-axis, based on the box size in the Z-dimension after equilibration. All systems were simulated using the CHARMM36m^45^ force field. The Charmm-modified TIP3P and Multisite Ca^2+^ parameters were used. We chose to use the CHARMM36m force field, rather than our Electronic Continuum Correction (ECC) version^34^ for two main reasons. Firstly our ECC version does not include compatible Ca^2+^ and NH_4_^+^ parameters and the same scaling factor of 0.78 is likely not realistic for these ions. Secondly we observe that under voltage the same binding sites are preferred, with higher overall occupancy when using ECC. In all simulations we used a 2 fs timestep and the Particle Mesh Ewald Method^46^ for the electrostatic interactions, using a cut-off distance of 1.2 nm. The force-switch method was used to turn Van der Waals interactions off from 1.0 to 1.2 nm. Semi-isotropic Parrinello-Rahman pressure coupling^47^ and the Nose-Hoover thermostat^48^ were used to keep the system at 1 bar and 310 K, respectively. Hydrogen involving bonds were constrained using the LINCS algorithm^49^. In the production runs with position restraints on backbone atoms force constants of 1000 kJ/mol/nm^2^ were used. Positions of ions in the SF were obtained using a custom FORTRAN program^13^. Atom densities in the cavity were calculated using a custom MDAnalysis-based^50,51^ python script. Permeation events were counted as previously described^34^.

### Protein expression and purification

A codon optimised DNA sequence encoding full length C-terminal His-tagged MthK was synthesised by GeneArt Life Technologies (Germany) cloned into the pET21a vector and transformed into *E. coli* BL21(DE3)pLysS.

In order to express perdeuterated, ^13^C- and ^15^N-labelled MthK, following a previously published deuterium adaptation protocol^38,52^, bacterial cultures were grown in perdeuterated M9 minimal media with ^15^ND_4_Cl and ^2^H^13^C labelled glucose (D-glucose-^13^C_6_,1,2,3,4,5,6,6-d_7_, Cambridge Isotope Laboratories, USA) as the sole nitrogen and carbon sources. For the production of protonated MthK, a protonated M9 medium with ^15^NH_4_Cl and ^13^C_6_-D_7_ glucose (Cambridge Isotope Laboratories, USA) as the sole nitrogen and carbon sources was used instead, and without a previous adaptation phase. Unlabelled MthK pore domain was expressed in a similar way using LB media.

In all cases, the protein was overexpressed for 16 hours at 25 °C after IPTG induction (0.5 mM) at an OD of 0.8. Cells were harvested, resuspended in lysis buffer [20 mM Tris (pH 8.0), 100 mM KCl, 1.4 mM ß-mercaptoethanol] and lysed using an LM10 microfluidizer (Microfluidics, USA) at 15,000 psi working pressure. Insoluble parts were removed by centrifugation, and the supernatant was incubated with 2% Decylmaltosid (w/w, DM, Glycon Germany) for 1 h at 4 °C. The solubilized protein was purified by cobalt-based gravity flow affinity chromatography (TALON® Superflow™, Cytiva, USA) with 0.2% DM (w/w) in wash [20 mM Tris (pH 8.0), 100 mM KCl, 10 mM Imidazole, 1.4 mM ß-mercaptoethanol] and elution buffer [20 mM Tris (pH 8.0), 100 mM KCl, 500 mM Imidazole, 1.4 mM ß-mercaptoethanol]. Purified full length MthK was digested with bovine pancreatic trypsin (Sigma-Aldrich) for 2 h at room temperature and the reaction was stopped by the addition of trypsin inhibitor type II (Sigma-Aldrich).

Isotope labelled, trypsin digested MthK (NMR samples) was mixed with asolectin (Sigma-Aldrich) in 5% DM (w/V) at a lipid to protein ration of 1:1 (w/w). Detergent was removed by dialysis against sample buffer (20 mM Tris (pH 8.0), 100 mM KCl, [or 100 mM ^15^NH_4_Cl] 1.4 mM ß-Mercaptoethanol) over the course of 10 days and buffer exchanges every second day. The sample was centrifuged for 2 h at 4 °C and 100,000 relative centrifugal force (rcf) and the pellet was used to fill MAS rotors by centrifugation in a benchtop centrifuge at 10,000 rcf.

Unlabelled, trypsin digested MthK (electrophysiology samples) was reconstituted into lipid vesicles composed of an asolectin phospholipid mixture (Avanti Polar Lipids). The asolectin phospholipid mixture was solubilized with 2% (w/V) DM detergent and dialysis (against 20 mM Tris-HCl, pH 7.6, 450 mM KCl) was used to slowly remove detergent from the detergent/lipid/protein mixture. Various protein-to-lipid ratios (0.1– 3 μg protein/mg lipid) were used in the reconstitution.

### Electrophysiology

Channel recordings were obtained using the Nanion Orbit Mini. Bilayers were painted with DPhPC (10 mg/ml) in decane using the ~100 μm MECA 4 chips filled with recording buffer (120 mM KCl, 80 mM KOH, 10 mM HEPES, 10 mM CaCl_2_, pH 7.6). The channel orientation was determined based on their inward rectification property. The traces were analyzed with ClampFit 11.2.

### Solid-state NMR

^13^C detected experiments were recorded on a 700 MHz Bruker spectrometer equipped with a 3.2 mm probe. A set of 3D experiments: (H)NCOCA, (H)NCACO, (H)CANCO, (H)NCACB and (H)NCO(CA)CB were recorded at 17 kHz magic-angle spinning (MAS) and at a sample temperature close to 0 °C. Heteronuclear magnetization transfers were achieved through cross-polarization (CP)^53^. Homonuclear (C-C) magnetization transfers were achieved through BSH-CP^54^ (for backbone CA-CO and CO-CA) and DREAM^55^ (for CA-CB). A detailed description of the procedure for acquiring these spectra is described in a previously published protocol^56^.

^1^H detected experiments were recorded on 600, 900 and 1200 MHz Bruker spectrometers equipped with 1.3 and 1.9 mm probes. Experiments for assignments were recorded on a sample with 100 mM ^15^NH_4_Cl and 100 mM CaCl_2_ at 900 MHz using a 4 channel (DHCN) 1.3 mm probe (Bruker Biospin) at 60 kHz MAS. A set of two double sensitivity-enhanced 3D spectra ((H)CANH and (H)CONH), and four triple sensitivity-enhanced 4D spectra [(H)CACONH, (H)COCANH, (H)CXCANH and (H)CXCA(CO)NH] were recorded for assignments, as previously described^37^. CP was used for H-C transfers, TROP for C-N transfers^57^, homoTROP for backbone C-C (CA-CO, CO-CA) transfers^58^ and DIPSI-3 for side-chain (CX-CA) transfers^59^. A deuterium lock was used to compensate for the drift of the magnetic field. Additional standard^52^ (at 40 kHz MAS, using a 1.9 mm probe) or sensitivity-enhanced^57^ (at 55-60 kHz MAS, using a 1.3 mm probe) 3D experiments were recorded on a 600 MHz spectrometer on the samples with 100 mM ^15^NH_4_ [(H)CANH, (H)CONH, (H)CA(CO)NH, (H)CO(CA)NH, (H)CB(CA)], 100 mM ^15^NH_4_ + 10 mM CaCl_2_ [(H)CANH, (H)CONH, (H)CA(CO)NH, (H)CO(CA)NH, (H)CB(CA)NH], 100 mM KCl [(H)CANH, (H)CONH] and 100 mM KCl + 10 mM CaCl_2_ [(H)CANH, (H)CONH] in order to transfer the assignments and compare the different samples.

Experiments for detection and assignments of the bound ^15^NH_4_ ions were performed as previously described^33,38^. INEPT was used for through-bond magnetization transfers with the INEPT delays set to 2.4-2.7 and 1.05-1.15 ms (depending on which spectrometer was used) after optimization around the theoretical values for N-H J-couplings (73.5 Hz) in ^15^NH_4_^60^. 2D (H)CH spectra with CP (7 ms mixing time) were used to characterize magnetization transfers between the H atoms of ^15^NH_4_ and backbone C atoms of MthK. Spin diffusion and RFDR^61^ (xy4^1^_4_ phase cycling) was used for H-H transfers in experiments that investigate ion-ion and ion-water interactions.

Spectra were processed using TopSpin 4 and nmrPipe^62^. Spectra recorded with non-uniform sampling (NUS) were reconstructed using compressed sensing with the iterative soft thresholding (IST) algorithm implemented in nmrPipe^63^ or the iterative re-weighted least squares (IRLS) algorithm in the qMDD software^64–66^. NUS lists were generated from http://gwagner.med.harvard.edu/intranet/hmsIST/gensched_new.html^67,68^. Analysis was performed using CCPNMR 3^69^.

## Acknowledgements

We thank Dagmar Michl for expert technical assistance and Prof. Han Sun for providing access to the Nanion Orbit Mini electrophysiology setup. This work was funded by the Leibniz-Forschungsinstitut für Molekulare Pharmakologie, the Leibniz Society within the Leibniz Collaborative Excellence funding programme (Project title: Ion Selectivity and Conduction Mechanism of Cation Channels, Project number: K305/2020 - to A.L., W.K., and B.L.d.G.) and the Deutsche Forschungsgemeinschaft (DFG, German Research Foundation)--under Germany’s Excellence Strategy--EXC 2008/1 (UniSysCat)--390540038 (to A.L.). C. Ö. acknowledges support from the Human Frontier Science Program (LT000303/2019-L).

## Author Contributions

Conceptualization: CÖ, RdV, WK, BLdG, AL

Methodology: CÖ, RdV, JL, CS

Software: RdV, WK, BLdG

Validation: CÖ, RdV, WK, BLdG, AL

Formal analysis: CÖ, RdV, JL, DQ

Investigation: CÖ, RdV, JL, DQ, CS, SL

Resources: BLdG, AL

Data curation: CÖ, RdV, WK, BLdG, AL

Writing - original draft: CÖ, RdV

Writing - review and editing: CÖ, RdV, WK, BLdG, AL

Visualization: CÖ, RdV

Supervision: WK, BLdG, AL

Project administration: CÖ, RdV, WK, BLdG, AL

Funding acquisition: CÖ, WK, BLdG, AL

## Data availability

Assigned chemical shifts have been deposited to the BMRB under accession numbers 53314 (^1^H detected data) and 53315 (^13^C detected data). The data that supports the findings of this study are available from the corresponding authors upon reasonable request.

