## Supplementary Information for "Atomistic Mechanism of Calcium-Mediated Inward Rectification of the MthK Potassium Channel by Solid-State NMR and MD Simulations"

(a) Research Unit Molecular Biophysics, Leibniz Forschungsinstitut für Molekulare Pharmakologie (FMP), Robert-Rössle-Straße 10, 13125 Berlin, Germany

(b) Computational Biomolecular Dynamics Group, Max Planck Institute for Multidisciplinary Sciences, Am Fassberg 11, 37077 Göttingen, Germany

(c) MOE Key Lab for Cellular Dynamics, School of Life Sciences, Division of Life Sciences and Medicine, University of Science and Technology of China, Hefei 230026, China

(d) Hefei National Research Center for Interdisciplinary Sciences at the Microscale, University of Science and Technology of China, Hefei, Anhui 230026, China

(e) Department of Chemistry, Queen Mary University of London, 327 Mile End Road, London E1 4NS, United Kingdom

(f) Institute of Biology, Humboldt-Universität zu Berlin, Invalidenstraße 42, 10115 Berlin, Germany.

#These authors contributed equally to this work.

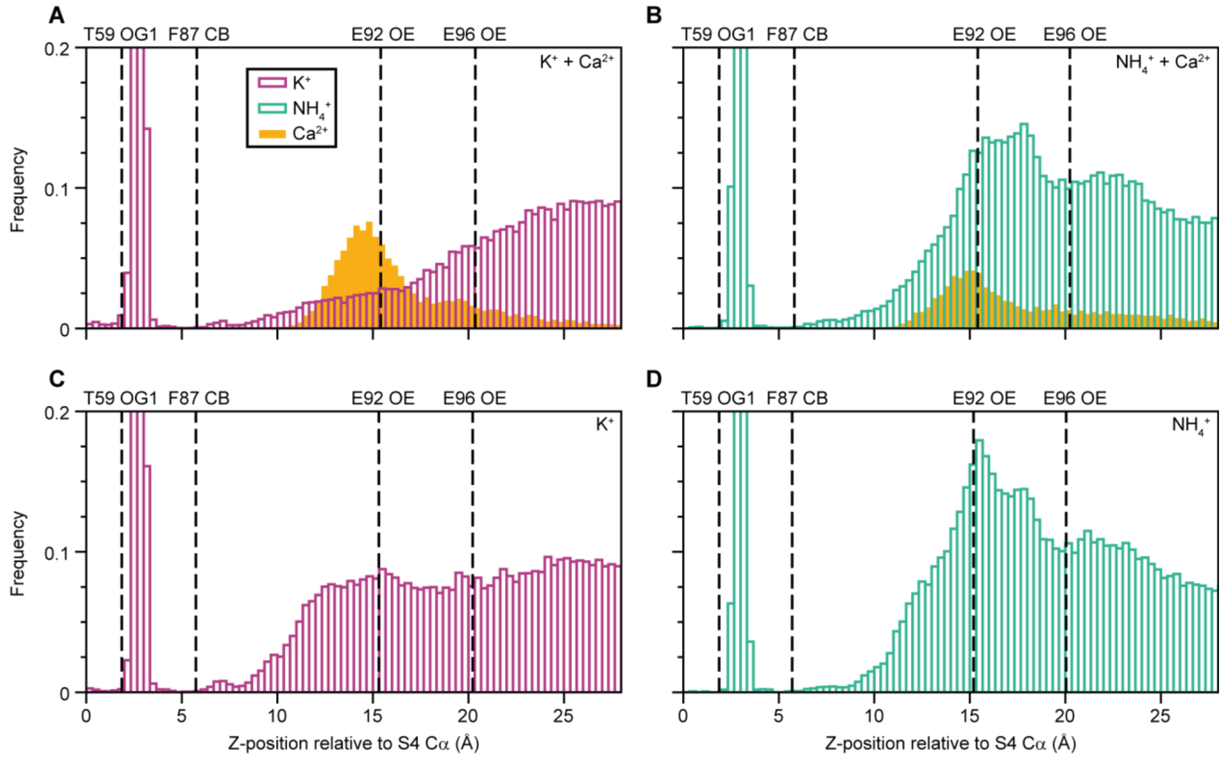


**Figure S1.** $Ca^{2+}$ binding in the MthK cavity under negative voltage. Ion densities in the cavity relative to T59 from MD simulations under -300 mV over 10 replicates (A) KCl + $CaCl_2$,(B) $NH_4Cl$ + $CaCl_2$,(C) KCl,(D) $NH_4Cl$.

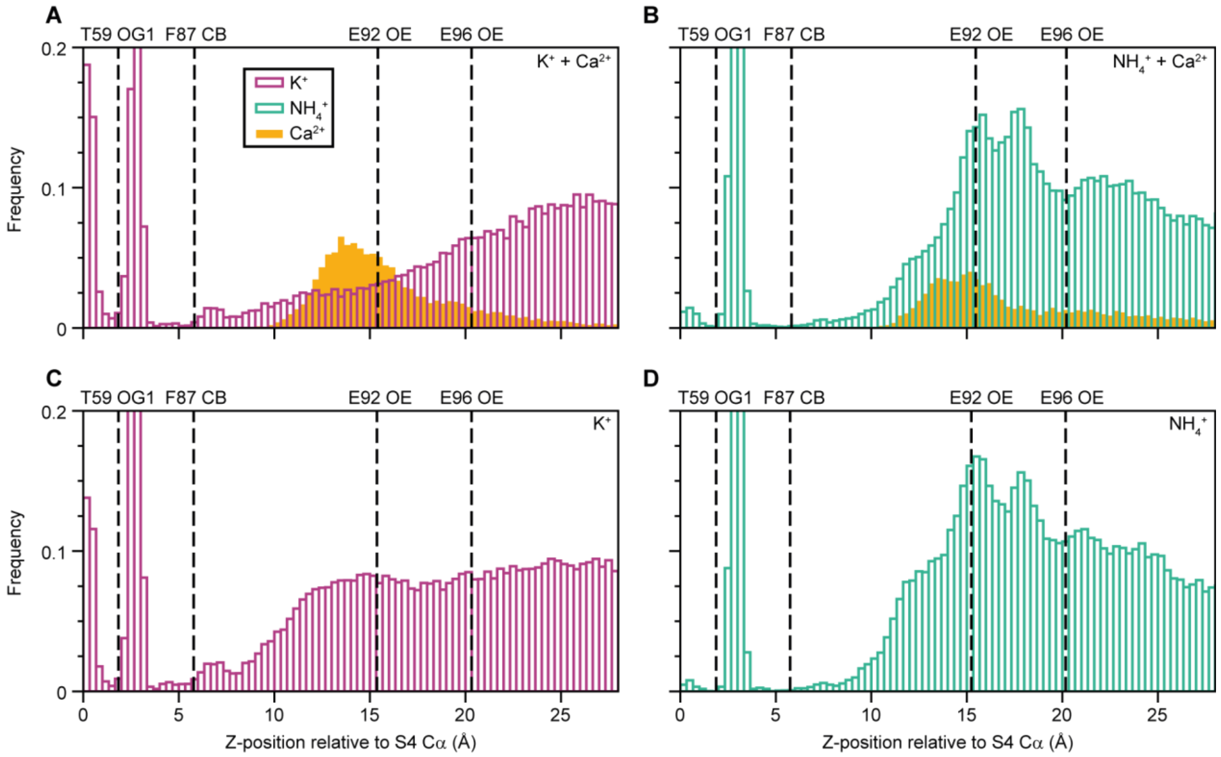


**Figure S2.** $Ca^{2+}$ binding in the MthK cavity without voltage. Ion densities in the cavity relative to T59 from MD simulations without voltage over 10 replicates (A) KCl + $CaCl_2$, (B) $NH_4Cl$ + $CaCl_2$,(C) KCl,(D) $NH_4Cl$.

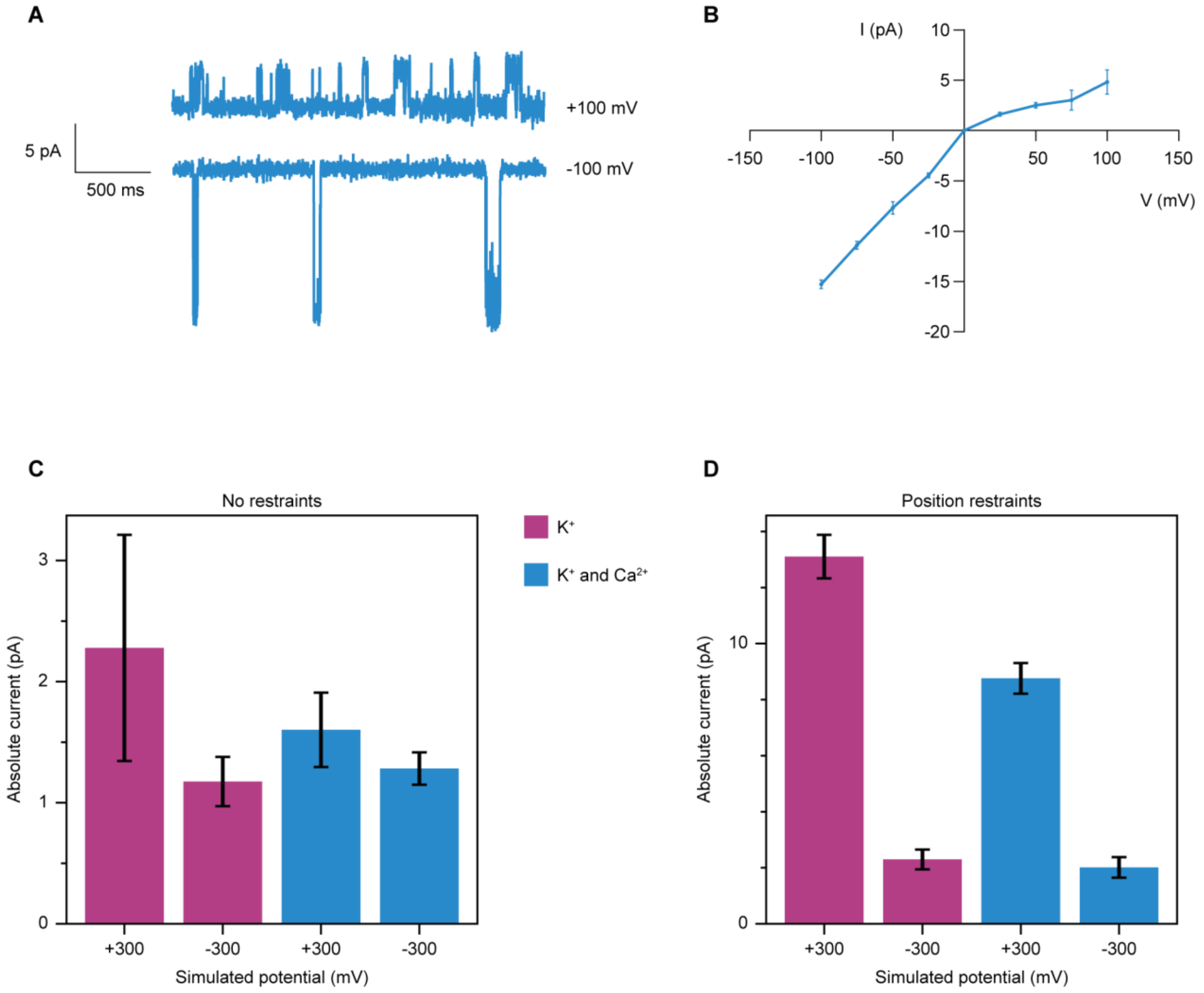


**Figure S3.** Experimental and simulated electrophysiology. (A) Examples of permeation events observed in electrophysiology experiments at +/- 100 mV. (B) I-V curve of MthK pore domain in the presence of 10 mM $Ca^{2+}$. Data points are mean and +/- standard error of the mean from multiple outbursts of one bilayer experiment. (C and D) Absolute simulated conductance with an applied voltage of +/- 300 mV with and without the addition of 30 mM $Ca^{2+}$. (C) Without restraints and (D) with position restraints applied to backbone atoms of residues 86-98. Error bars in (C) and (D) represent standard error of the mean over 10 independent replicates.

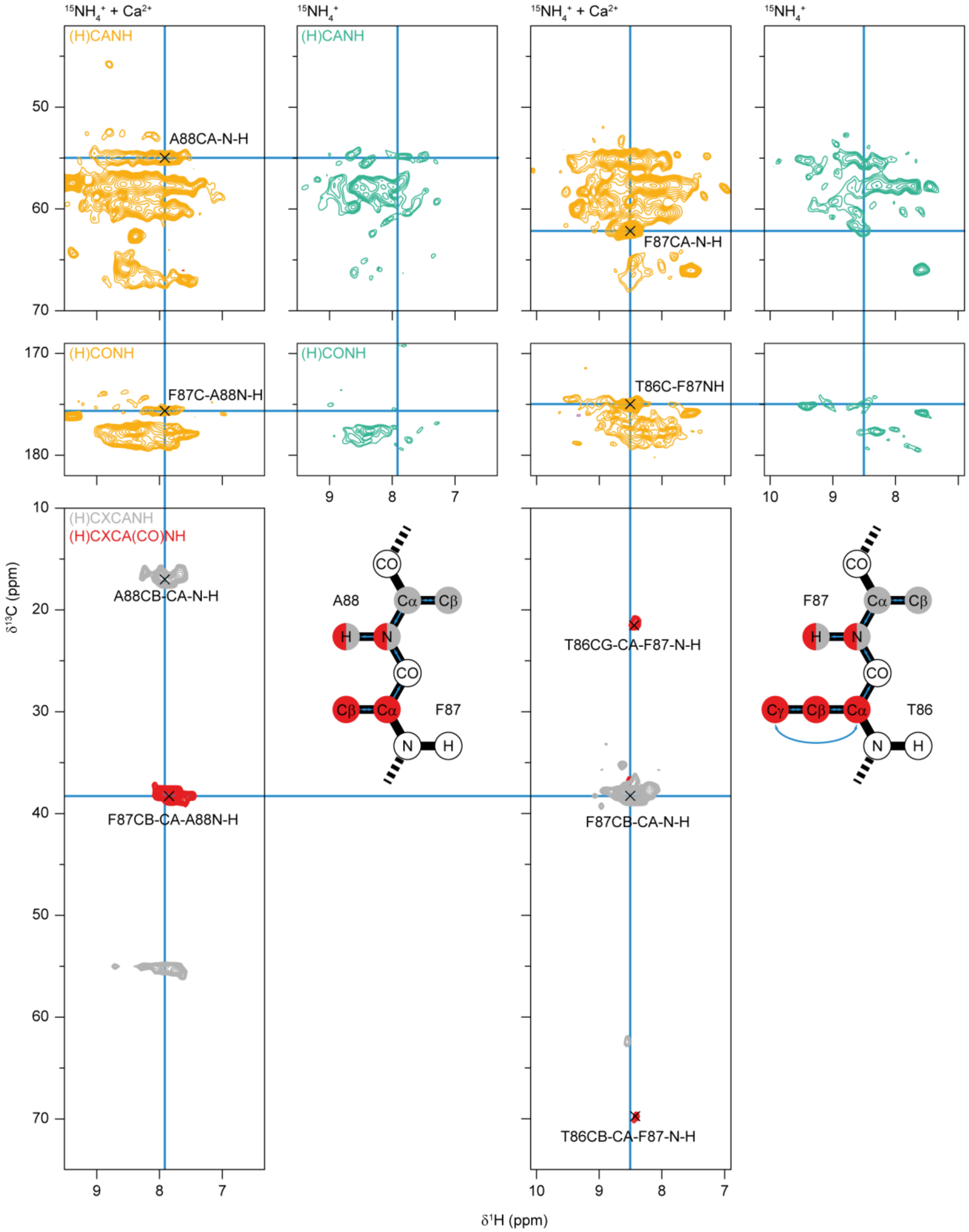


**Figure S4.** Example strip plots for the assignment of residues around the F87 $Ca^{2+}$ binding site. Top, 2D CH strip plots of (H)CANH and (H)CONH (orange for the sample with 100 mM $^{15}NH_4^+$ and 10 mM $Ca^{2+}$, green for the sample with only $^{15}NH_4^+$) taken at the $^{15}N$ chemical shift of A88 (left) and F87 (right). Bottom, 2D CXH strip plots of 4D (H)CXCANH (grey) and (H)CXCA(CO)NH recorded on a sample with 100 mM $^{15}NH_4^+$ and 100 mM $Ca^{2+}$. The CXH strips of the (H)CXCANH are taken at the $^{15}N$ and $^{13}CA$ chemical shifts of residue i (A88 - left and F87 - right), while the CXH strip plots of the (H)CXCA(CO)NH are taken at the $^{15}N$ chemical shift of residue i (A88 - left, F87 - right) and the $^{13}CA$ chemical shift of residue i-1 (F87 - left, T86 - right). See also the schematic drawings indicating the atoms for which the peaks are observed in the 4D spectra.

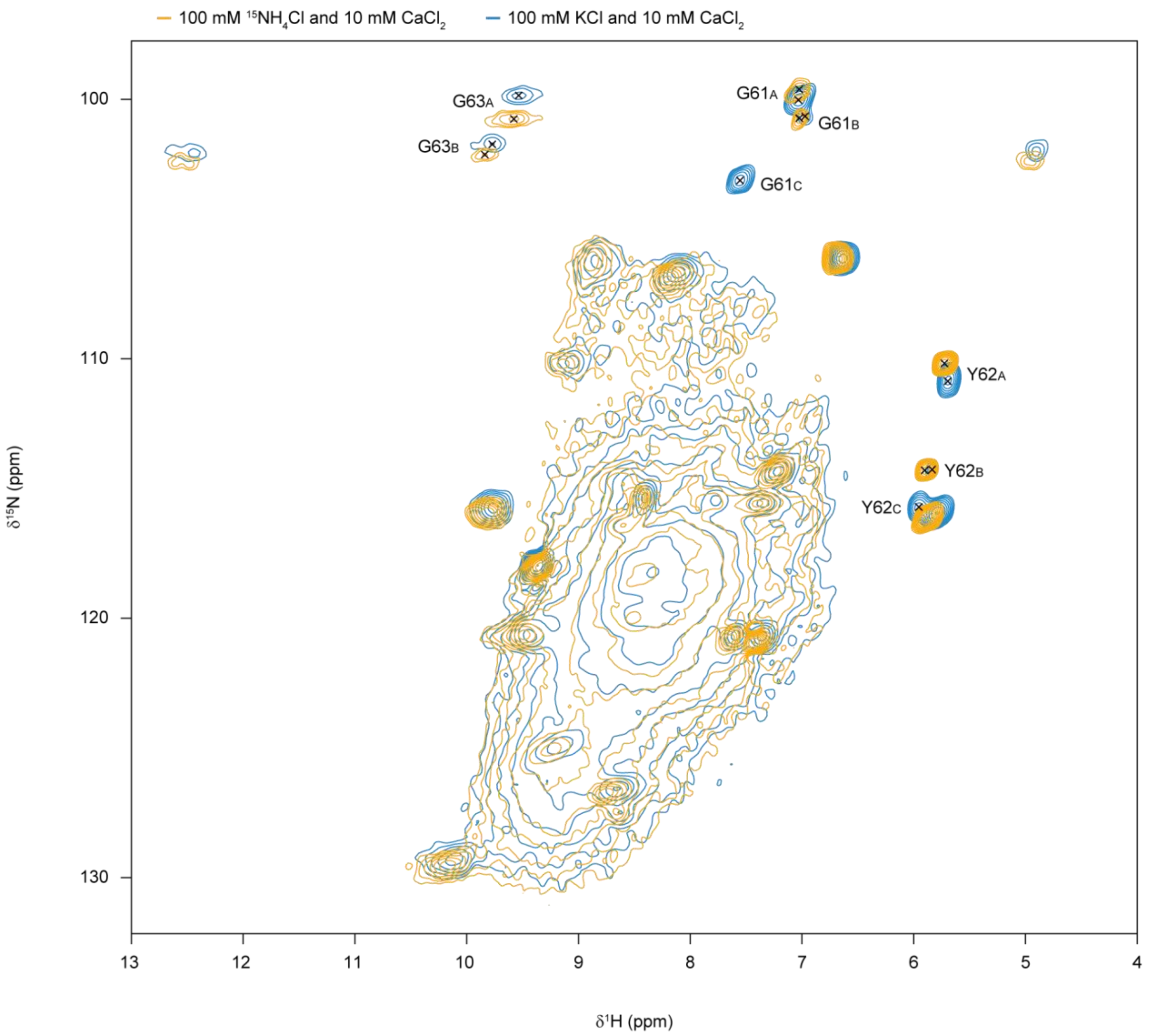


**Figure S5.** Comparison of samples with $K^+$ and $^{15}NH_4^+$. 2D (H)NH spectra of MthK with $^{15}NH_4^+$ (orange) and $K^+$ ions (blue). Both samples contain 10 mM $Ca^{2+}$ ions. The different conformations of the selectivity filter residues are labelled (with "A","B","C").

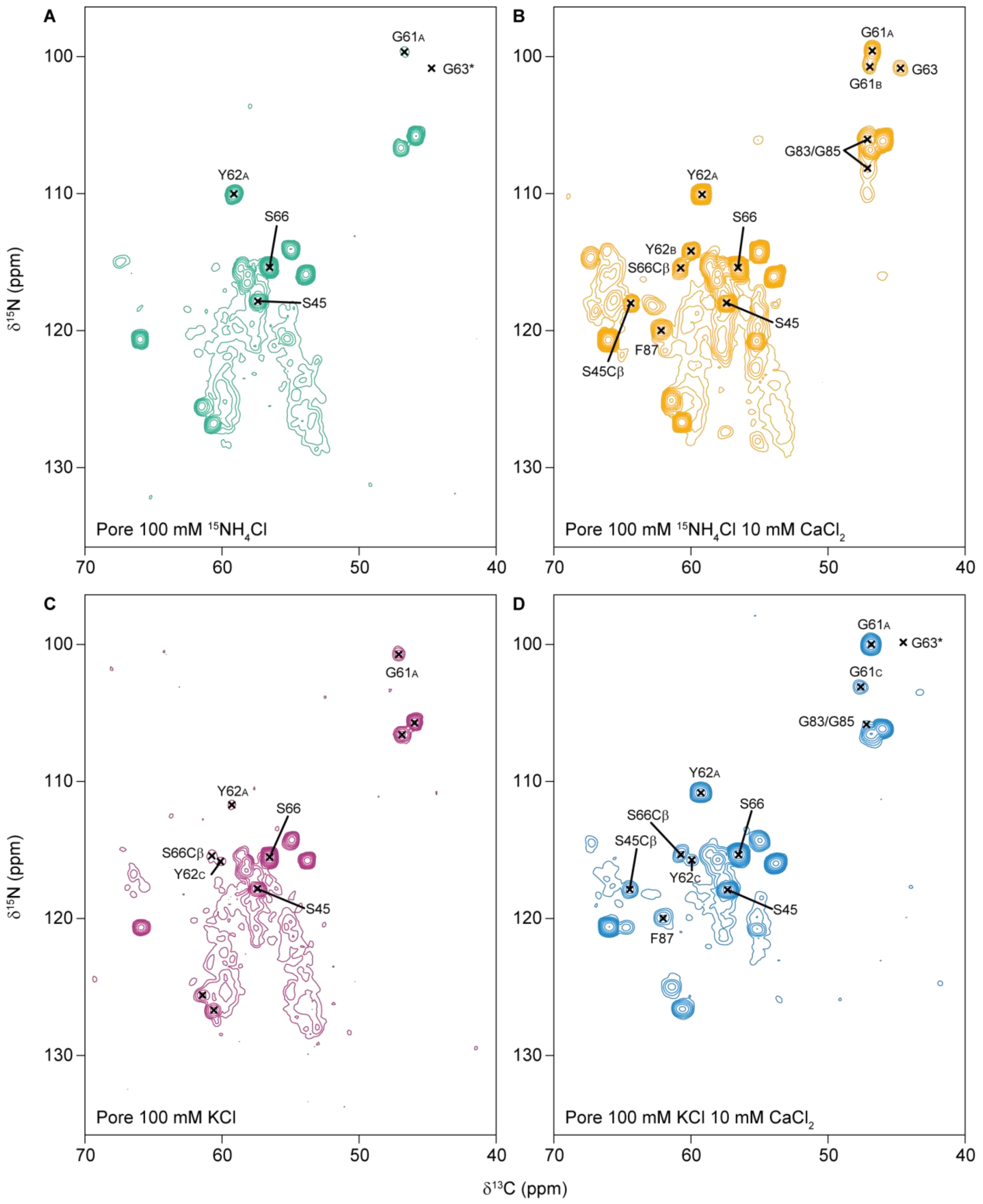


**Figure S6.** 2D NCA projections from $^{1}H$ detected 3D (H)CANH spectra of MthK pore domain with 100 mM $^{15}NH_4Cl$ (A, green), 100 mM $^{15}NH_4Cl$ + 10 mM $CaCl_2$ (B, orange), 100 mM KCl (C, purple), and 100 mM KCl + 10 mM $CaCl_2$ (D, blue). Peaks from the selectivity filter residues and other residues that are affected by $Ca^{2+}$ are labelled. This includes serine N-CB peaks, that appear with high intensity in the presence of $Ca^{2+}$ ions, and residues around the F87 $Ca^{2+}$ ion binding site.

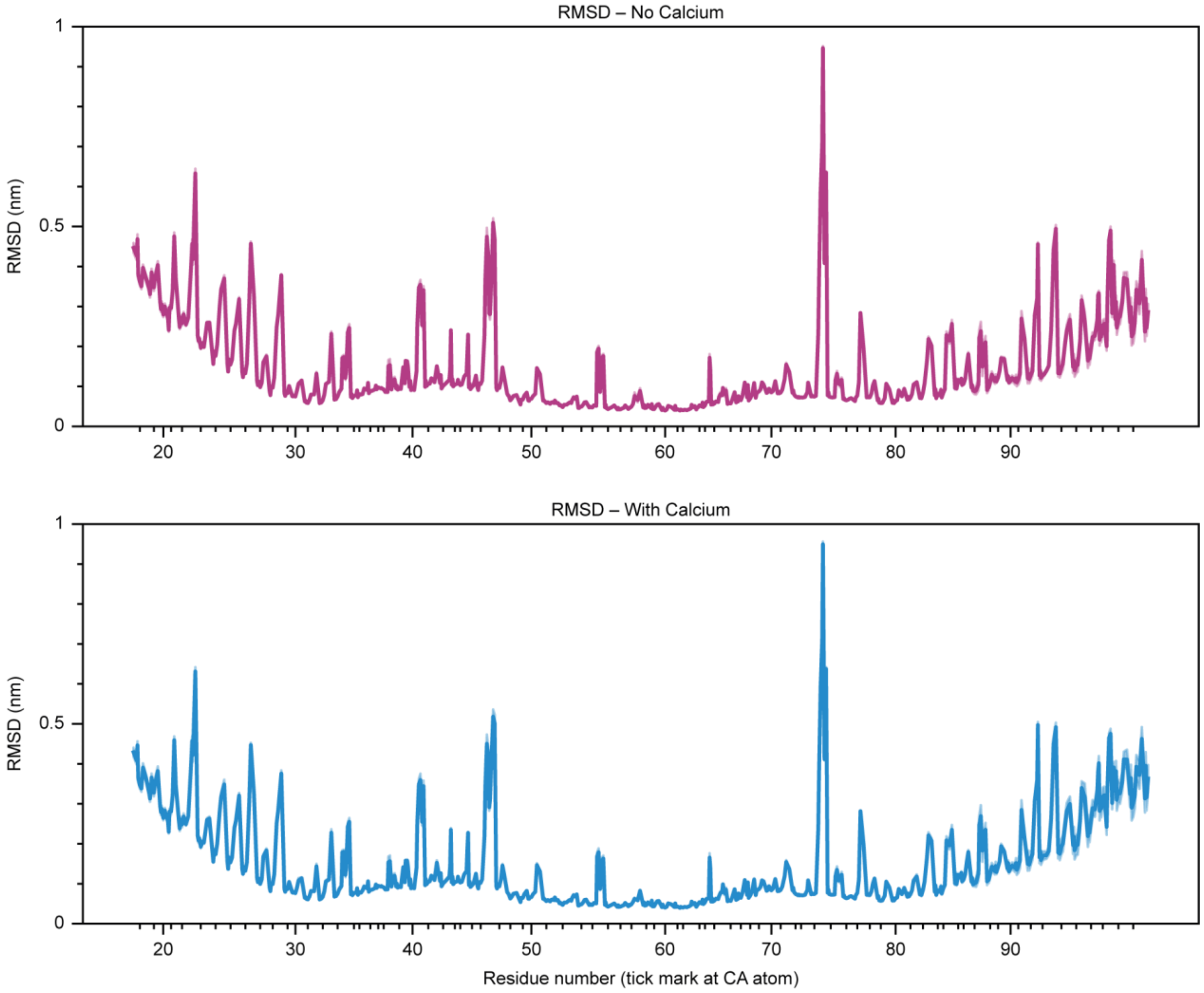


**Figure S7.** RMSD plots for MD simulations of MthK with $K^+$, without (top, purple) and with (bottom, blue) $Ca^{2+}$.

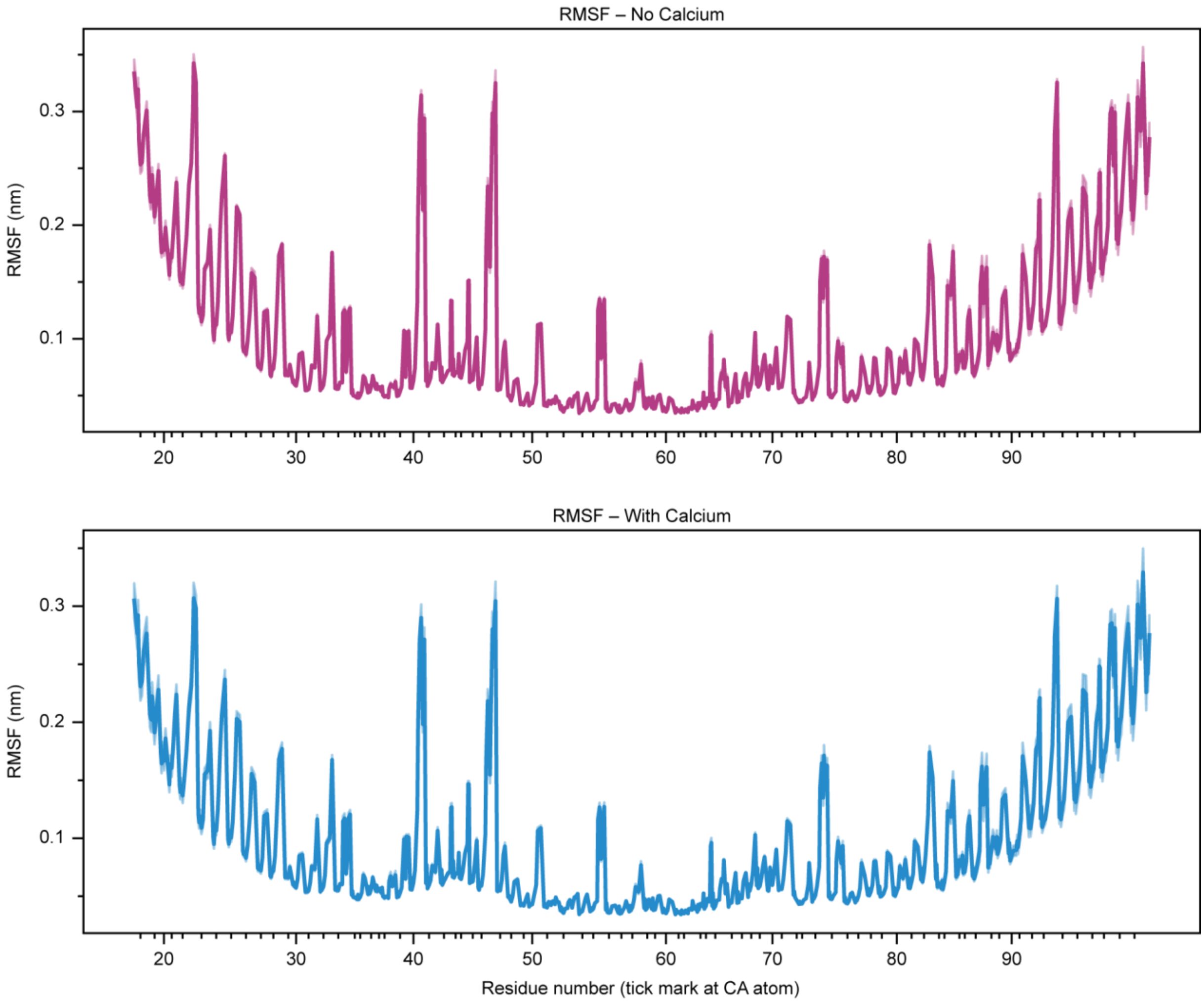


**Figure S8.** RMSF plots for MD simulations of MthK with $K^+$, without (top, purple) and with (bottom, blue) $Ca^{2+}$.

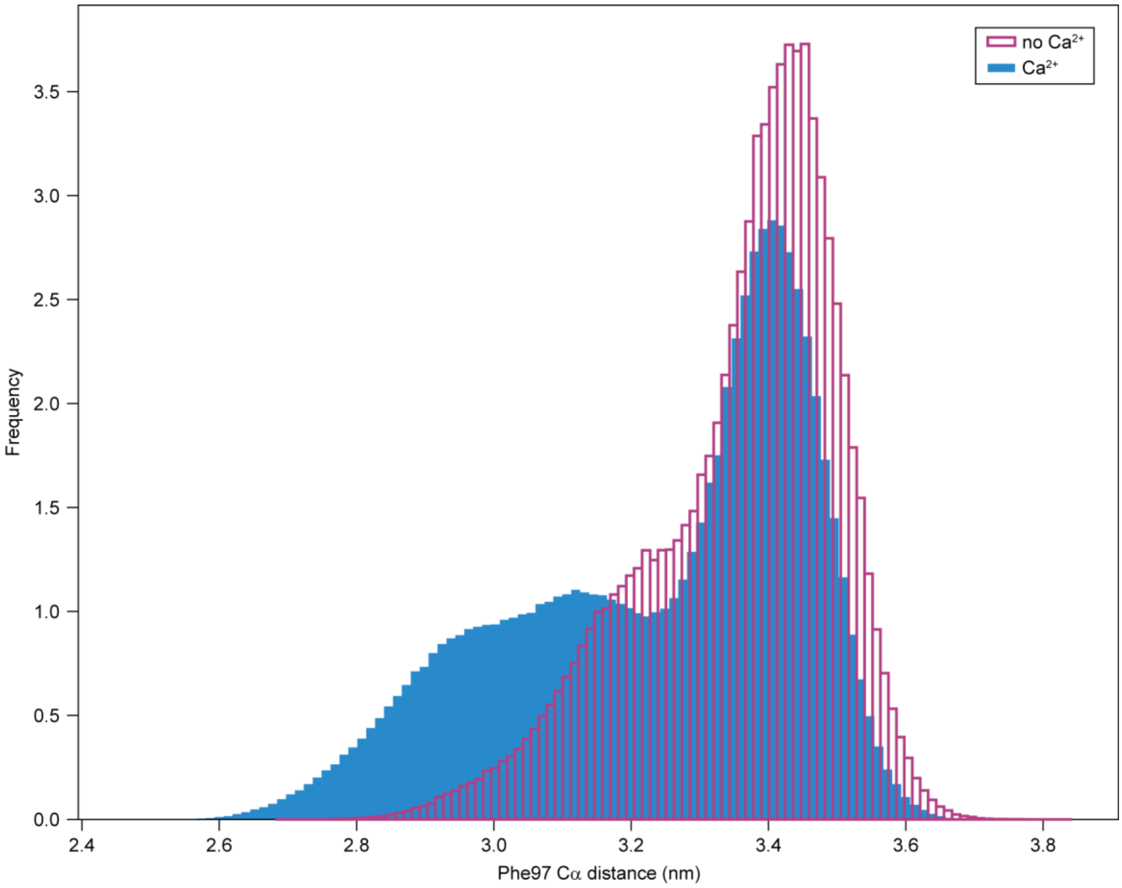


**Figure S9.** Lower gating opening distances. Mean distance between opposing F97 Cα atoms. Averaged over 50 5 μs simulations starting from 5 different initial opening distances.

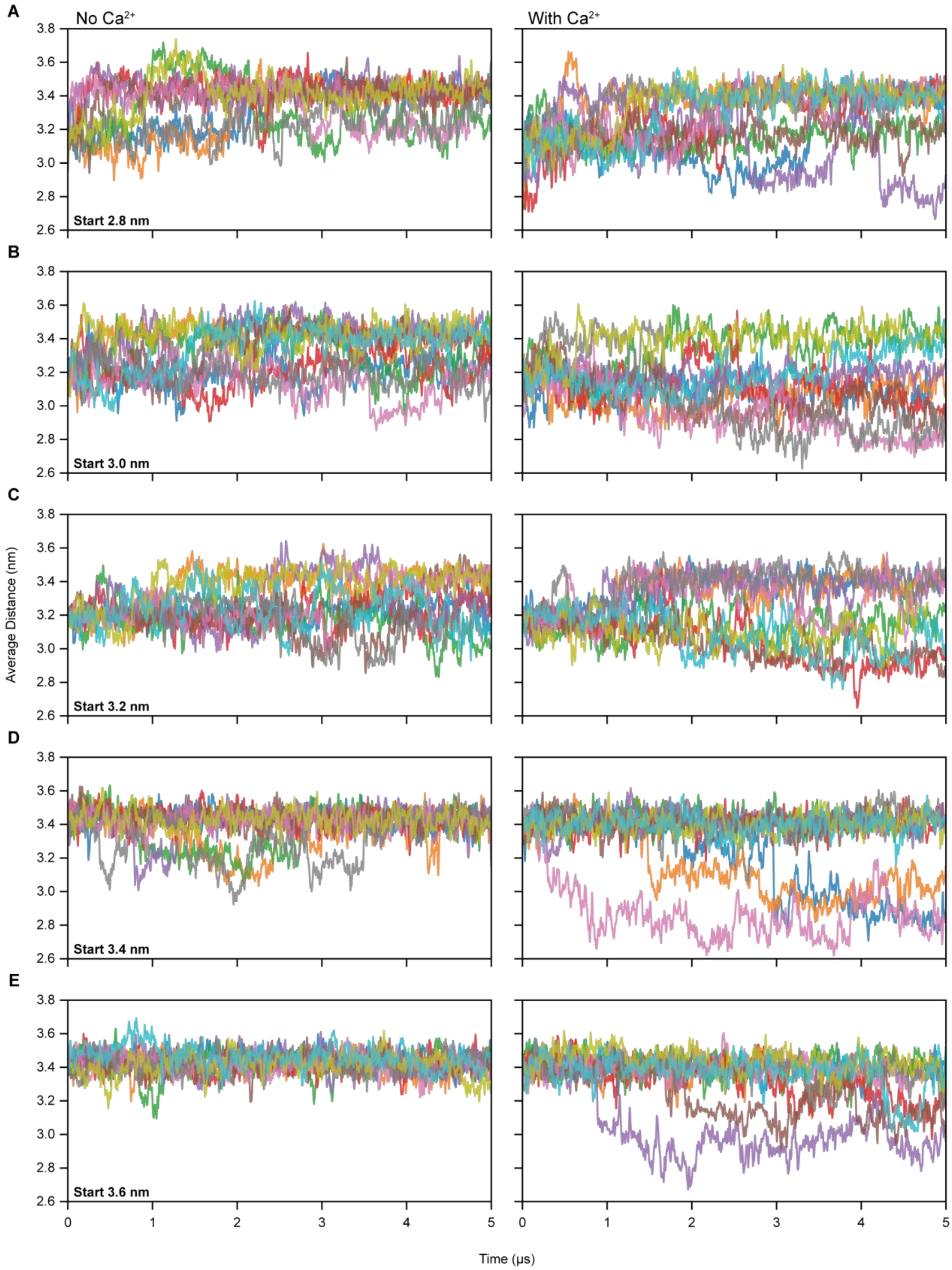


**Figure S10.** Smoothened Timetraces of the average F97 Cα distance between opposing subunits. Each subplot shows traces from a specific starting condition based on the initial F97 Cα distance and whether calcium is included.

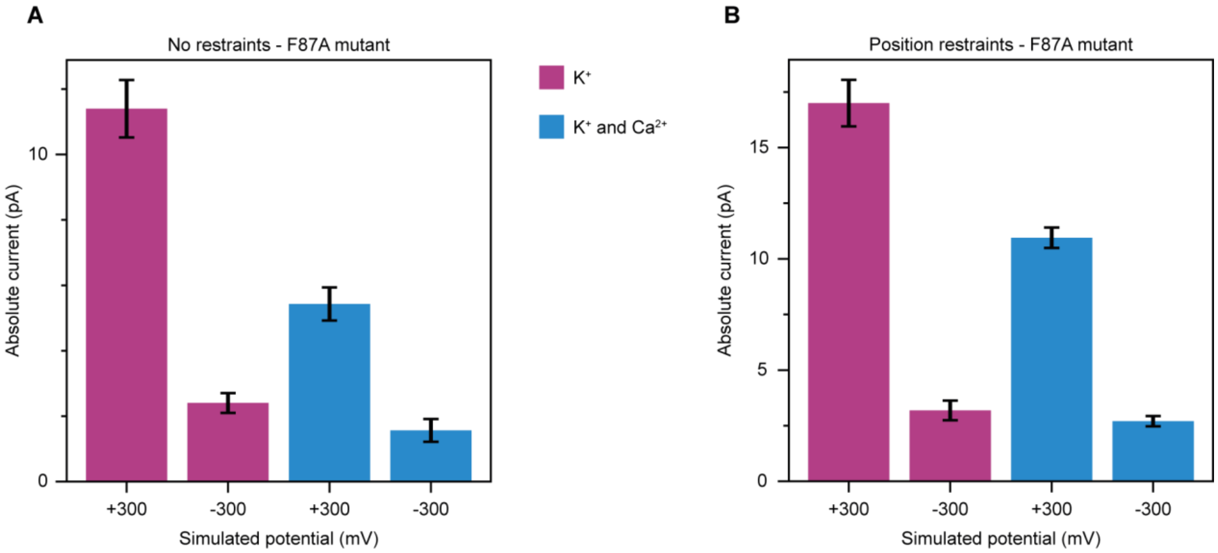


**Figure S11.** Simulated electrophysiology of MthK F87A. (A and B) Absolute simulated conductance with an applied voltage of +/- 300 mV with and without the addition of 30 mM $Ca^{2+}$. (A) Without restraints and (B) with position restraints applied to backbone atoms of residues 86-98. Error bars in (A) and (B) represent standard error of the mean over 10 independent replicates.

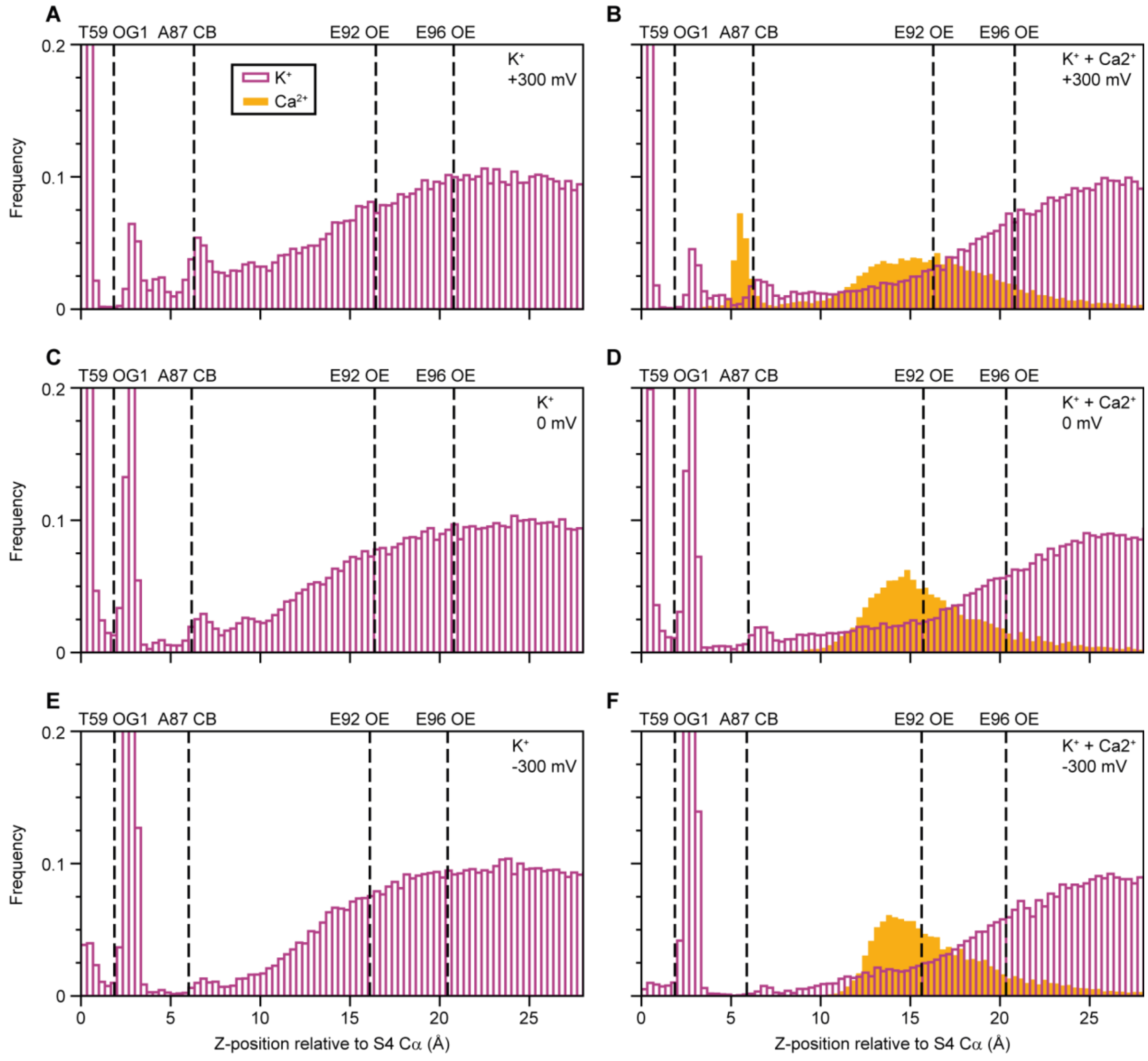


**Figure S12.** Ion densities in the MthK F87A cavity relative to Thr59 from MD simulations under different voltages with and without $Ca^{2+}$ (A) KCl at 300 mV. (B) KCl + $CaCl_2$ at 300 mV. (C) KCl at 0 mV. (D) KCl + $CaCl_2$ at 0 mV. (E) KCl at -300 mV. (F) KCl + $CaCl_2$ at -300 mV.

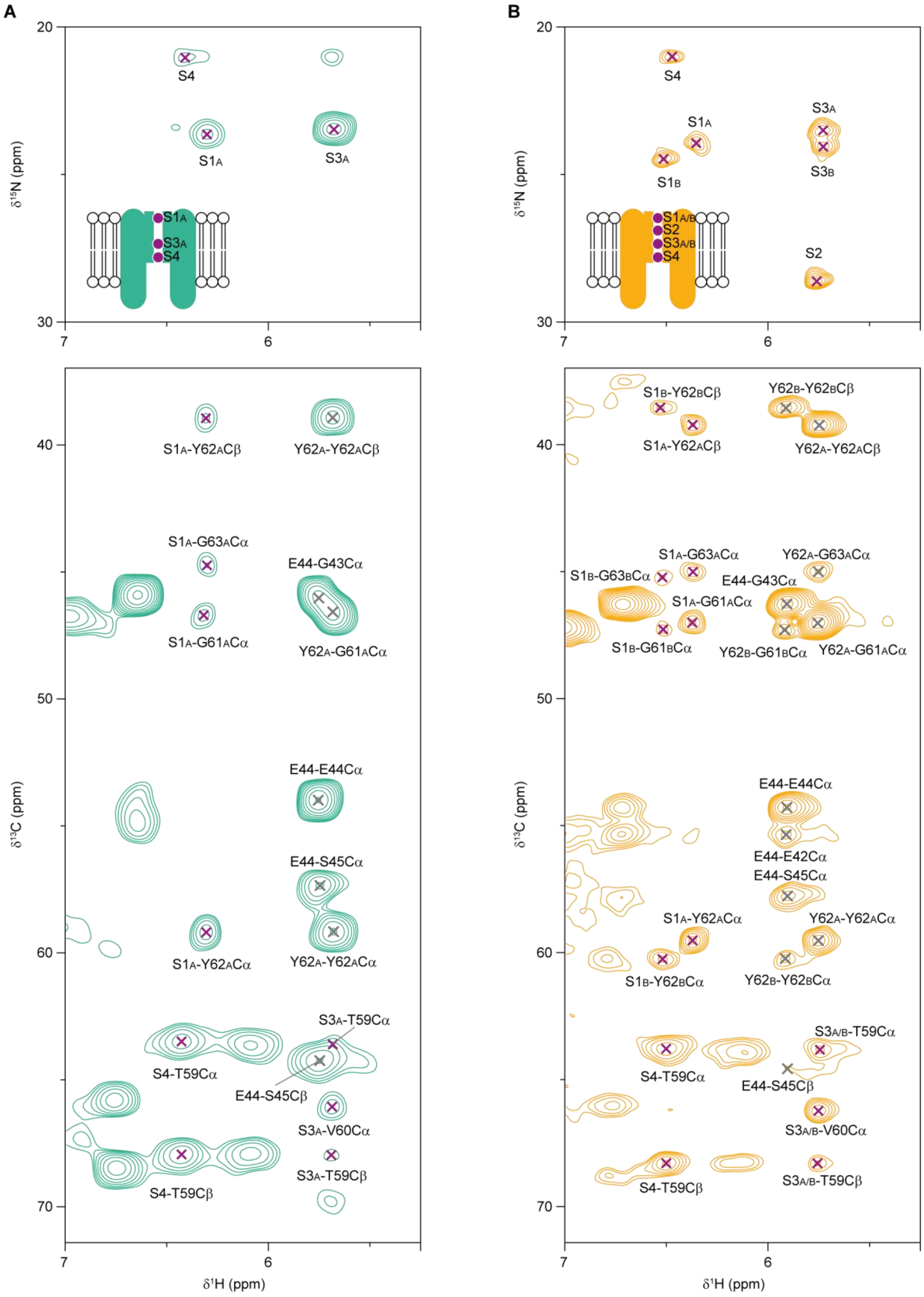


**Figure S13.** Detection of bound $^{15}NH_4^+$ ions in the MthK pore domain. $^1H$ detected INEPT-based 2D (H)NH spectra of $^{15}NH_4^+$ (top) and CP-based (H)CXH spectra (bottom) recorded on $^2H^{13}C^{15}N$ labelled MthK pore domain samples with 100 mM $^{15}NH_4Cl$ (A, green spectra) and 100 mM $^{15}NH_4Cl$ + 10 mM $CaCl_2$ (B, orange spectra). Peaks involving $^{15}NH_4^+$ ions are labelled with purple crosses and peaks between backbone atoms are labelled with dark grey crosses.

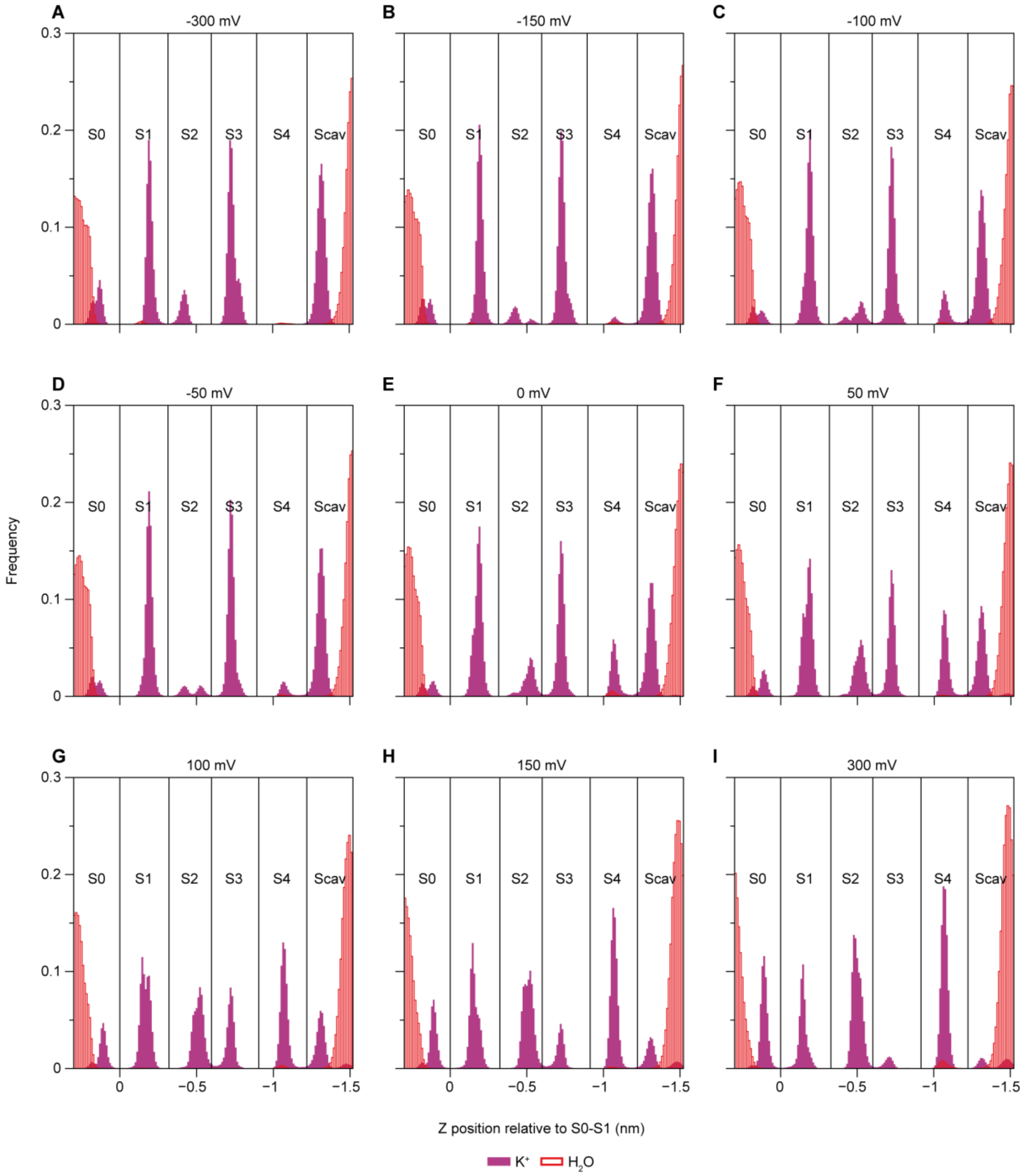


**Figure S14.** Influence of voltage on SF occupancy for simulations with $K^+$ ions. (A to I) $K^+$ and water occupancies at -300 to +300 mV, in steps of 50 mV. $K^+$ ions are labelled in purple and water in red.

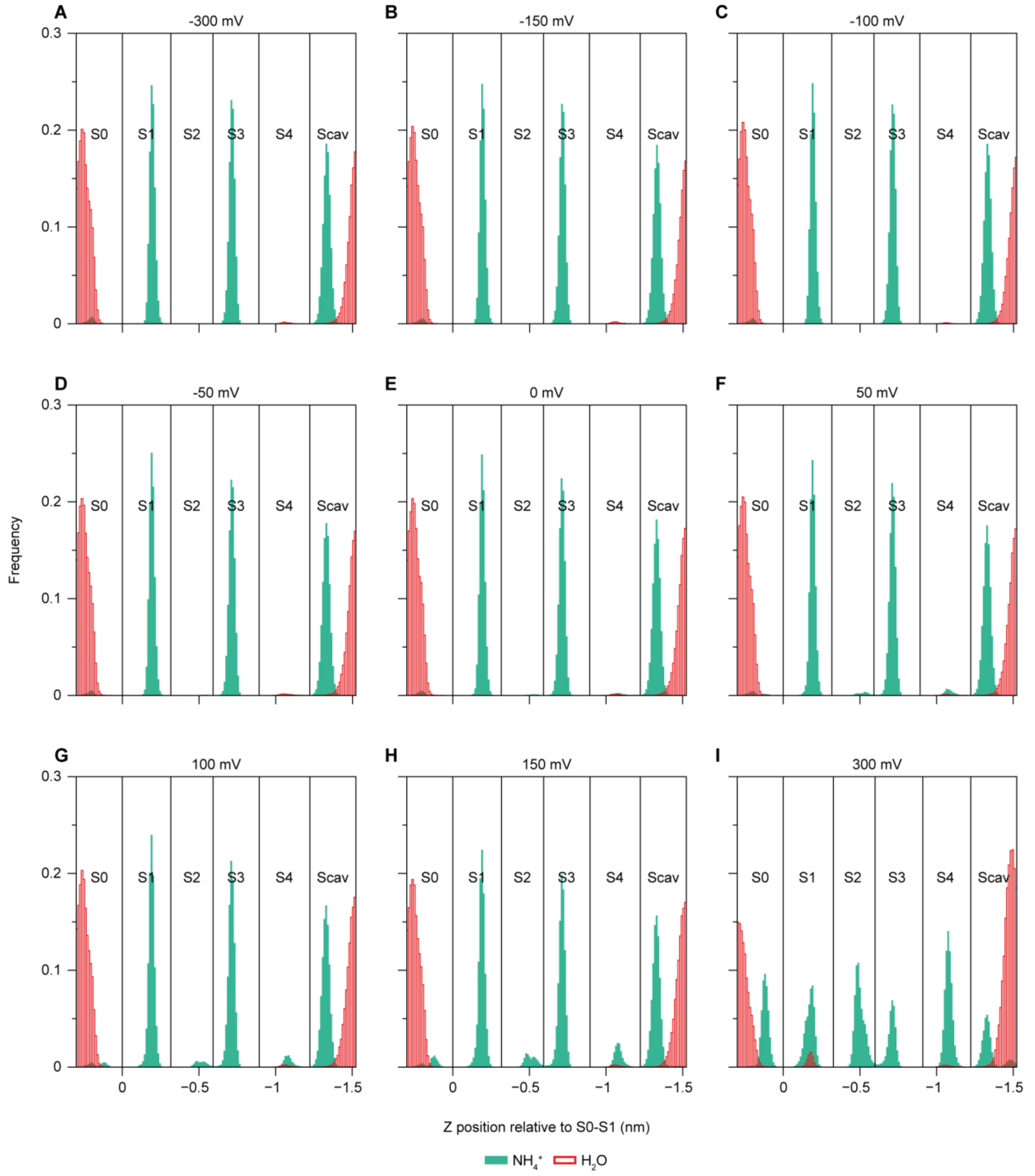


**Figure S15.** Influence of voltage on SF occupancy for simulations with $NH_4^+$ ions. (A to I) $NH_4^+$ and water occupancies at -300 to +300 mV, in steps of 50 mV. $NH_4^+$ ions are labelled in green and water in red.

**Table S1.** Chemical shift assignments (deposited in the BMRB under ID: 53315) based on $^{13}C$ detected experiments on $^{13}C^{15}N$ labelled MthK pore domain with 100 mM KCl. Values are in ppm and are referenced externally using DSS.

| Residue number | Residue type | N | C | CA | CB |
|---|---|---|---|---|---|
| 28 | LEU | - | 177.99 | 58.15 | - |
| 29 | ALA | 120.39 | 178.92 | 55.62 | - |
| 30 | VAL | 116.65 | 177.45 | 66.87 | 31.40 |
| 31 | ILE | 119.46 | 179.75 | 64.37 | 37.54 |
| 32 | ILE | 123.55 | 176.32 | 66.30 | 37.77 |
| 33 | TYR | 120.54 | 177.99 | 61.51 | 39.59 |
| 34 | GLY | 103.64 | 174.94 | 48.42 | |
| 35 | THR | 116.98 | 176.75 | 67.41 | 68.98 |
| 36 | ALA | 121.02 | 179.38 | 55.12 | 18.08 |
| 37 | GLY | 104.62 | 174.29 | 47.56 | |
| 38 | PHE | 123.92 | 177.84 | 63.39 | 38.61 |
| 39 | HIS | 115.72 | 178.09 | 58.10 | 26.54 |
| 40 | PHE | 115.12 | 176.85 | 60.89 | |
| 41 | ILE | 117.11 | 176.83 | 64.53 | 38.08 |
| 42 | GLU | 114.62 | 177.40 | 55.21 | |
| 43 | GLY | 106.16 | 174.69 | 46.32 | |
| 44 | GLU | 115.34 | 175.92 | 53.53 | 30.60 |
| 45 | SER | 118.66 | 175.45 | 57.52 | 64.93 |
| 46 | TRP | 126.38 | 177.89 | 61.84 | 28.43 |
| 47 | THR | 115.81 | 175.86 | 67.70 | 69.48 |
| 48 | VAL | 120.97 | 177.67 | 66.27 | 31.10 |
| 49 | SER | 115.97 | 176.49 | 63.51 | |
| 50 | LEU | 128.24 | 178.74 | 57.70 | 42.15 |
| 51 | TYR | 122.09 | 176.16 | 60.67 | 38.48 |
| 52 | TRP | 121.36 | 178.94 | 61.91 | 27.70 |
| 53 | THR | 121.36 | 176.66 | 68.83 | |
| 54 | PHE | 119.16 | 177.16 | 63.88 | 39.75 |
| 55 | VAL | 117.80 | 178.11 | 66.36 | 32.10 |
| 56 | THR | 119.94 | 175.99 | 68.03 | |
| 57 | ILE | 114.90 | 175.23 | 64.97 | 35.95 |
| 60 | VAL | 122.05 | 177.28 | 66.15 | 31.58 |
| 61 | GLY | 101.16 | 173.28 | 47.30 | |
| 62 | TYR | 112.08 | 178.08 | 59.82 | 39.84 |
| 63 | GLY | 100.15 | 175.17 | 44.80 | |
| 64 | ASP | 120.69 | 175.48 | 55.42 | 36.25 |
| 65 | TYR | 115.41 | 174.34 | 57.32 | 41.38 |
| 66 | SER | 115.67 | 169.69 | 56.66 | 61.00 |
| 67 | PRO | | 176.21 | 62.48 | |
| 68 | SER | 116.17 | 174.52 | 58.41 | 64.88 |
| 69 | THR | 115.30 | 172.79 | 58.56 | 70.26 |
| 70 | PRO | | 178.27 | 66.04 | |
| 71 | LEU | 117.24 | 178.77 | 58.39 | 41.20 |
| 72 | GLY | 106.60 | 177.98 | 47.17 | |
| 73 | MET | 127.40 | 177.62 | 61.05 | 31.09 |
| 74 | TYR | 118.41 | 178.91 | 63.46 | 38.59 |
| 75 | PHE | 119.84 | 178.87 | 60.21 | 38.05 |
| 76 | THR | 122.12 | 175.67 | 68.71 | 67.74 |
| 77 | VAL | 121.71 | 177.26 | 68.45 | 30.25 |
| 78 | THR | 108.43 | 174.78 | 66.05 | 69.09 |
| 79 | LEU | 123.37 | 178.66 | 57.43 | 41.95 |

**Table S2.** Chemical shift assignments (deposited in the BMRB under ID: 53314) based on $^{1}H$ detected experiments on 100% $H_2O$ back-exchanged $^{2}H^{13}C^{15}N$ labelled MthK pore domain with 100 mM $^{15}NH_4Cl$ and 10 or 100 mM $CaCl_2$. Values are in ppm and are referenced externally using DSS.

| Residue number | Residue type | H | N | C | CA | CB | CG |
|---|---|---|---|---|---|---|---|
| 38 | PHE | | | 177.88 | 63.05 | 38.24 | |
| 39 | HIS | 7.81 | 115.69 | 177.93 | 57.87 | 26.73 | |
| 41 | ILE | | | 176.83 | 64.17 | | |
| 42 | GLU | 7.24 | 114.35 | 177.66 | 54.99 | 27.56 | 35.73 |
| 43 | GLY | 6.67 | 106.59 | 174.79 | 46.04 | | |
| 44 | GLU | 5.77 | 116.06 | 176.02 | 53.72 | 29.75 | 34.18 |
| 45 | SER | 9.37 | 117.92 | 175.67 | 57.38 | 64.67 | |
| 46 | TRP | 9.16 | 124.88 | 177.77 | 61.51 | 28.14 | |
| 47 | THR | 8.60 | 114.83 | 175.72 | 67.42 | 69.29 | 20.47 |
| 48 | VAL | 7.62 | 120.67 | 177.54 | 66.05 | 30.13 | 22.48 |
| 49 | SER | | | 176.27 | 63.23 | | |
| 50 | LEU | 8.63 | 127.68 | | 57.53 | 41.35 | |
| 59 | THR | | | 171.75 | 63.55 | 67.96 | |
| 60A | VAL | | | 176.63 | 66.09 | | |
| 61A | GLY | 6.96 | 99.63 | 173.28 | 46.75 | | |
| 62A | TYR | 5.66 | 110.01 | 178.63 | 59.18 | 38.87 | |
| 63 | GLY | 9.59 | 100.86 | 175.22 | 44.64 | | |
| 64 | ASP | 9.43 | 120.99 | 175.58 | 55.11 | 35.77 | |
| 65 | TYR | 7.32 | 115.76 | 174.19 | 57.37 | 40.27 | |
| 66 | SER | 8.37 | 115.18 | 169.82 | 56.60 | 60.84 | |
| 67 | PRO | | | 175.82 | 62.33 | 31.50 | |
| 68 | SER | 9.75 | 115.73 | 174.65 | 58.00 | 64.71 | |
| 69 | THR | 8.58 | 115.54 | 173.21 | 58.57 | | 20.99 |
| 70 | PRO | | | 178.24 | 65.84 | | |
| 71 | LEU | 8.70 | 116.51 | 178.57 | 58.16 | 40.22 | |
| 72 | GLY | 8.13 | 106.61 | 177.87 | 46.86 | | |
| 73 | MET | 8.69 | 126.67 | 177.66 | 60.72 | 30.13 | 32.34 |
| 74 | TYR | 8.34 | 118.11 | 178.75 | 63.08 | 37.57 | |
| 75 | PHE | 9.28 | 119.74 | | 60.03 | 38.14 | |
| 86 | THR | | | 175.16 | 66.15 | 69.76 | 21.00 |
| 87 | PHE | 8.48 | 120.21 | 175.55 | 62.11 | 38.22 | |
| 88 | ALA | 7.81 | 118.71 | 180.66 | 55.05 | 16.94 | |
| 89 | VAL | 7.77 | 116.68 | 177.16 | 64.71 | 33.66 | |
| 60B | VAL | | | 175.63 | | | |
| 61B | GLY | 6.97 | 101.06 | 174.38 | 46.83 | | |
| 62B | TYR | 5.83 | 114.27 | 178.53 | 60.01 | 38.21 | |

**Table S3.** Chemical shift assignments of the different conformations of the selectivity filter residues for MthK pore domain with $K^+$ ions. Assignments are based on $^1H$ detected (H)NH, (H)CANH, and (H)CONH spectra recorded on 100% $H_2O$ back-exchanged $^2H^{13}C^{15}N$ labelled MthK pore domain with 100 mM KCl and 10 mM $CaCl_2$. Values are in ppm and are referenced externally using DSS.

| Residue number | Residue type | H | N | C | CA |
|---|---|---|---|---|---|
| 60A | VAL | | | 177.02 | |
| 61A | GLY | 7.01 | 100.01 | 173.26 | 46.88 |
| 62A | TYR | 5.68 | 110.83 | 178.21 | 59.28 |
| 63A | GLY | 9.55 | 99.78 | | 44.64 |
| 60B | VAL | | | 175.58 | |
| 61B | GLY | 6.96 | 100.66 | 174.47 | |
| 62B | TYR | 5.83 | 114.22 | | |
| 63B | GLY | 9.77 | 101.72 | | |
| 60C | VAL | | | 179.29 | |
| 61C | GLY | 7.55 | 103.10 | 174.60 | 47.63 |
| 62C | TYR | 5.94 | 115.68 | | 59.94 |

**Table S4.** Chemical shift assignments of $^{15}NH_4^+$ ions bound in the selectivity filter of MthK pore domain. Assignments are based on a 2D (H)NH INEPT spectrum and 2D (H)COH and (H)CXH CP spectra recorded on a sample with 100 mM $^{15}NH_4Cl$ and 10 mM $CaCl_2$. Values are in ppm and are referenced externally using DSS.

| Ion binding site | H | N |
|---|---|---|
| S1A | 6.36 | 23.94 |
| S1B | 6.51 | 24.48 |
| S2 | 5.76 | 28.61 |
| S3A | 5.74 | 23.53 |
| S3A | 5.73 | 24.06 |
| S4 | 6.47 | 21.03 |
